# Riding the savory horse: An active mindset and food macronutrient composition influence attentional bias toward food cues

**DOI:** 10.1101/2025.06.05.658045

**Authors:** Marc Ballestero-Arnau, Borja Rodríguez-Herreros, Manuel Moreno-Sánchez, Toni Cunillera

## Abstract

Food cues that appear in the visual field capture our attention easily and can influence eating behavior. The current study investigated the influence of food-related stimuli on visual attention, considering the macronutrient composition of food items. Images representing sweet and savory foods were employed, the latter consisting primarily of high-protein foods. The participants were primed with these images prior to performing the attentional task. We found that both sets of food images elicited an emotional attentional blink (EAB), but a stronger EAB was observed for the high-protein foods, and this observation was further supported by a negative correlation between the attentional bias (ABias) and the proportion of protein consumed by the participants before the experiment, with participants who consumed less protein exhibiting a stronger ABias toward high-protein foods. These findings suggest that an ABias might also arise to facilitate the consumption of high-protein foods when prior consumption of this macronutrient is low.

## INTRODUCTION

In recent decades, the promotion and accessibility of ultra-processed, calorically dense foods have undergone an unprecedented surge in modern societies. This dietary shift may be partially explained by the pervasive influence of neuromarketing strategies aimed at enhancing the appeal and consumption of these foods (Hill & Peters, 1998; Lieberman, 2006). This prevailing dietary style poses a challenge to the homeostatic mechanisms responsible for body weight regulation (Berthoud, 2012), which has culminated in a steadily increasing prevalence of obesity and associated comorbidities that have reached pandemic proportions (Di Cesare et al., 2016; WHO, 2016).

At the cognitive level, attention seems to play a fundamental role in the development and maintenance of eating habits (Johnson, 2013; Rangel, 2013). Numerous studies have provided evidence that food stimuli—particularly palatable foods—capture visual attention more effectively than nonfood-neutral cues do (Nijs et al., 2010). Additionally, factors such as caloric content are believed to further enhance this effect (Cunningham & Egeth, 2018a). Analogous to evidence from emotional stimuli (Kennedy & Most, 2015; Most et al., 2005), food appears to receive priority in attentional processing, a phenomenon that may be particularly salient in individuals with obesity (Castellanos et al., 2009; Yokum et al., 2011). Pictures of calorically dense foods can automatically capture attention, even when they are completely irrelevant to the task (Ballestero-Arnau et al., 2021; Cunningham & Egeth, 2018a; Kennedy et al., 2024). Similarly, food stimuli can interfere with an ongoing goal, inducing food consumption even when the energy balance is stable (Berthoud et al., 2007; Johnson, 2013). Individuals who are susceptible to overeating exhibit attentional bias (ABias) toward high-calorie foods in a situation of food deprivation (Castellanos et al., 2009; Dobson & Dozois, 2004). Accordingly, ABias toward food-related cues are considered a cognitive determinant underlying overeating in individuals with obesity (Field et al., 2016). Nonetheless, existing research is inconclusive, with two recent meta-analyses suggesting a lack of association between body mass index (BMI) and ABias (Hagan et al., 2020; Hardman et al., 2021).

The inconsistency regarding the association between ABias and obesity could be attributed to the oversight of within-individual fluctuations in ABias (Werthmann et al., 2016). Werthmann et al. induced either healthy or palatable temporary mindsets to investigate their effects on ABias toward food-related stimuli. Their results highlighted the significant influence of state-specific mindsets on attention toward food, which was beyond the influence of being a restrained eater. Additionally, other studies have shown that food-related thoughts held in working memory can influence attentional allocation (Higgs, Robinson, et al., 2012; Higgs, Rutters, et al., 2012; Rutters et al., 2015), highlighting the putative impact of mindset variations not only on food perception but also on attentional modulation toward food cues.

The temporal dimension of the ABias toward food has been studied using the rapid serial visual presentation (RSVP) task (Kirsten et al., 2019; Neimeijer et al., 2013a; Piech et al., 2010). In the classic RSVP paradigm, participants are instructed to detect two predefined targets embedded in streams where stimuli are displayed at a rapid rate of approximately 100 msec, and the time between the two targets (i.e., the lag) is manipulated. Originally designed to investigate goal-driven attentional limitations toward stimuli based on predetermined goals (Theeuwes, 2019), this task consistently reveals that accuracy in identifying a second target decreases when a previous target is presented in close temporal proximity. This temporal limitation of attention is a robust phenomenon known as attentional blink (Dux & Rentmarois, 2009; Raymond et al., 1992; Shapiro et al., 1997). An interesting variant of the classical RSVP task, known as the emotional attentional blink (EAB), has also been successfully used to investigate how food items capture attention (Arumäe et al., 2019; Ballestero-Arnau et al., 2021; Davidson et al., 2018; Neal et al., 2023; Piech et al., 2010). However, the EAB effect is consistently weaker than that of standard attentional blink (Kirsten et al., 2019; Neimeijer et al., 2013a; Santacroce et al., 2023). In the EAB task, participants are instructed to detect a single target in the RSVP stream. A similar attentional effect is thought to occur when performance is disrupted with a distractor image placed right before the target compared with when it is placed eight or nine images before (e.g., Lag2/Lag3 vs. Lag8/Lag9). Such stimulus-driven emotional capture has been reported in numerous single-target RSVP experiments (e.g., (Arnell et al., 2007; Ciesielski et al., 2010; Most et al., 2005); see (McHugo et al., 2013) for a review).

ABias is sensitive to changes in the motivational value of food, which is concurrently enhanced in situations of food deprivation. It has been proposed that cognitive and metabolic functions interact to create the motivational drive, reflected as a feeling of hunger, that leads to consummatory eating behaviors with the sole purpose of restoring one’s energy balance (Cunningham & Egeth, 2018b; Kaisari et al., 2019; Y. Liu et al., 2019; Neimeijer et al., 2013b). This motivational drive originates with the creation of reward expectancies from unspecific highly energetic food cues (Berridge, 2009a; Johnson, 2013; Zheng & Berthoud, 2008). Food-related stimuli could thus capture attention more easily in a situation of hunger, inciting one to act accordingly (Lang et al., 1997; Seibt et al., 2007). Such ABias has been related to the incentive salience of high-caloric food cues (Berridge, 2009b), especially with food stimuli representing high fat and/or sugar content (Greenberg & St. Peter, 2021; Simpson & Raubenheimer, 2014a). However, it is still unclear whether that motivational drive may simply urge the search for energy or otherwise underly the search for specific macronutrients, exerting an additional adaptive physiological function (Simpson & Raubenheimer, 2014b).

In the present study, we investigated two possible determinants of the modulation of ABias toward food: i) an active mindset to facilitate the attentional capture of food stimuli and ii) the consideration of attention as a basic cognitive function that could promote opportunistic behaviors to obtain specific macronutrients. We addressed the assumed specificity of the ABias toward high-energy food by comparing the attentional effect elicited by food images with high sugar and/or fat content, representative of rapid access to an energy supply, with food images characterized by elevated protein content. We induced an EAB effect by prompting two different mindsets with food images, matched for calories, that served as distractors in a single-target version of the RSVP task. Furthermore, participants were deceived to take part in a memory task to remember as many items as possible from a menu flyer within 30 s. Afterward, they were instructed to complete a task for which the memorized food items were completely irrelevant—the single-target version of the RSVP task—aimed at postponing the recall of the previously memorized items.

## METHODS EXPERIMENT 1

### Participants

We recruited 105 students from the Faculty of Psychology at the University of Barcelona to participate in the study. We administered the experiment in an online format. To avoid including spurious data, we excluded a participant if any of the following criteria were met: i) the mean duration of image presentation within the series was greater than 120 ms; ii) reaction times were greater than 2.5 standard deviations (S.D.) from the mean; iii) participants’ performance in conditions without the food distractor was less than 2.5 S.D. from the mean; and iv) a participant reported following a diet that excluded certain types of food (e.g., vegetarian, vegan). Data from 9 participants were excluded in accordance with these prestipulated exclusion criteria. Thus, the final sample was composed of 96 participants, who were randomly assigned to the sweet (46) and savory (50) food conditions when they were included in the experiment (Table 1). The two samples had similar sex, age, and BMI distributions (all p values > 0.5). All of the participants provided informed consent before they participated in this study, and they were compensated with course credits for completing the experiment. The study was approved by the local ethics committee.

**Table 1.** Demographic and anthropometric characteristics of the participants in Experiments 1 and 2 by food-type condition (sweet vs. savory) and food-reward-assignment condition (sweet vs. savory), respectively.

|  | Experiment 1 |  | Experiment 2 |  |
| --- | --- | --- | --- | --- |
|  | Sweet | Savory | Sweet | Savory |
| Sample size | 46 | 50 | 60 | 57 |
| Females | 41 | 43 | 54 | 45 |
| Males | 5 | 7 | 6 | 12 |
| Age | 19.5 (1.8) | 19.4 (1.9) | 19.7 (1.9) | 19.9 (1.8) |
| BMI-mean | 21.5 (2.7) | 21.9 (2.6) | 21.5 (2.7) | 22.5 (3.2) |
| BMI-range | 16.0-29.7 | 17.6-28.0 | 16.9-29.0 | 16.7-30.5 |

**Note.** The values are presented as the means with standard deviations in parentheses for age and BMI and as absolute counts for sample size and sex. The BMI range indicates the minimum and maximum body mass index observed within each group.

### Stimuli and procedure

We used a total of 37 images as visual stimuli during the experiment. All of the images were obtained from the Food-Pics database (Blechert et al., 2019). The pictures were arranged into three different categories. The first category included pictures of sweet foods with a high sugar content (ice cream, chocolate cookies, donuts, etc.). The second category was named savory. Food images belonging to this class were discernable by their high protein content (burgers, salamis, hot dogs, etc.). Finally, the last category included nonfood images from different thematic categories (Figure 1). We first conducted a set of statistical analyses to ensure that, apart from the sugar and protein contents, the perceptual and energetic properties of the two sets of foods were similar to each other (see the Supplementary Material).

**Figure 1.**
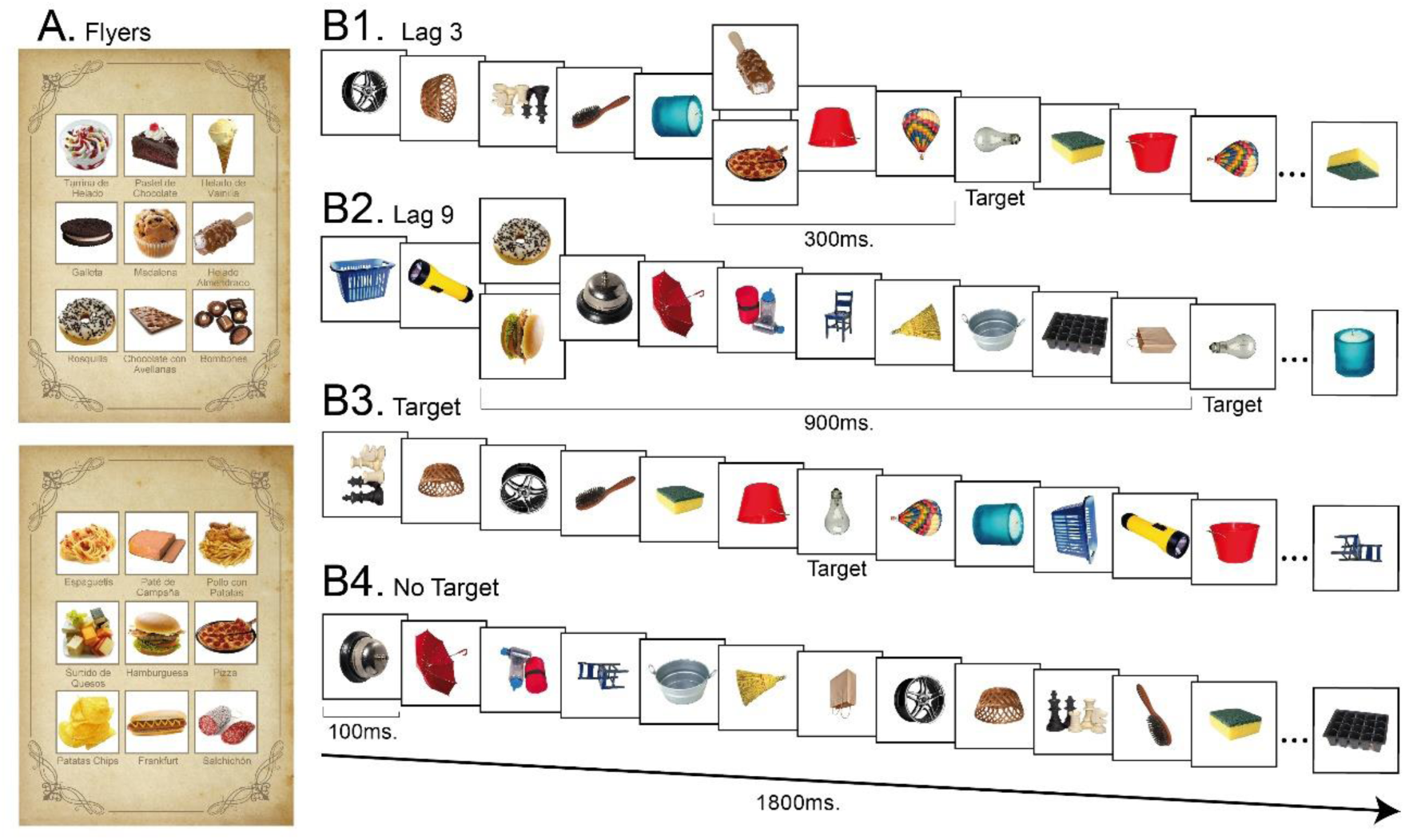
Illustration of the flyers used for the memory task and an example of a series in the RSVP task for each condition in Experiment 1. **A.** Flyers created in the “sweet” (upper) and “savory” (bottom) group conditions. **B1**. Series from Lag-3 condition; the distractor appeared 300 msec before the target (light bulb). **B2.** Series from the Lag-9 condition; the distractor was presented 900 msec before the target. **B3.** Series in which the target was presented without the distractor. **B4.** Series in which neither the target nor the distractors were presented. All series were composed of 18 images with a duration of 1800 msec. During the task, all distractor images appeared 12 times, six times in each Lag condition, distributed equiprobably in different positions along the image sequences. Note that the two distractor types are illustrated simultaneously, but in the real task, only one type of distractor (either a sweet or savory item) was presented across the task.

We programmed and executed the task as an online experiment using the PsyToolkit method for running online experiments (Stoet, 2017). For the experiment, each participant was randomly assigned to one of the two experimental food-type conditions automatically when they entered the website. Next, demographic data were collected, and instructions on how to proceed were presented on the screen. The first task consisted of a simple memory task. The participants were instructed to remember as many items as possible from a flyer presented on the screen for 30 s. The flyer contained either 9 pictures of sweet or savory foods, depending on the food-type condition. Immediately after the memory task was completed, the RSVP task was proposed as an ordinary exercise to delay memory recall, and participants were informed that after completing it, they would be asked to report as many foods as they could recall from the abovementioned flyer. The RSVP task was a version of a previous EAB paradigm (Most et al., 2005), with the requirement of detecting a single target in each trial (i.e., a picture of a light bulb, Figure 1). The task consisted of 6 blocks of 72 trials. In each trial, a series of 18 images were presented for 100 ms each (SOA = 0 ms; duration of a series = 1800 ms). The participants were asked to report at the end of each sequence, using the mouse buttons (left=yes; right=no), whether they had seen the target or not. The target was presented in 50% of the trials, appearing between the 6^th^ and 16^th^ positions within the sequence of images. In half of the trials where the target stimulus was presented, a distractor image was included. The distractor was one of the images from the flyer. Each of the 9 pictures that participants were asked to memorize in the preceding memory task appeared 12 times during the RSVP task in a random order and were equally distributed within the sequences. Crucially, the distractor images were placed within or outside the temporal window in which the attentional blink is known to occur (Schwabe et al., 2011), i.e., 300 ms prior to onset (Lag-3 condition) and 900 ms before the onset of the target (Lag-9 condition). To avoid a possible effect of stimulus habituation, all images presented along the series were randomly rotated at 0°, 90°, 180° and 270°.

We collected several subjective measurements to control for potential confounding factors. Before completing the memory task, the participants were asked to provide their weight and height and to indicate their subjective hunger level on a horizontal 100-point line presented on the center of the screen via a computer mouse, with 0 signifying *no hunger at all* and 100 indicating *very hungry*. Furthermore, after completing the RSVP task, the participants were asked to type the responses to the following questions: i) all of the items they could recall from the memory task (*memory recall*) and ii) the images from the flyer that they realized to have seen during the RSVP task (*distractor salience*). They were then instructed to provide a detailed account of their most recent meal, including all foods and beverages consumed. Finally, other subjective measurements were needed: iii) how much they liked the food presented in the flyer (*explicit food liking*), iv) how much they would have liked to consume those foods at that moment (*explicit food craving*), v) the perceived caloric content of each food item (*perceived calories*), and vi) the frequency of consumption of those foods (*consumption frequency)*. The participants provided their subjective rating using a similar horizontal 100-point line for each one, except for the consumption frequency, for which they used a 5-point Likert scale, with the option *rarely*, *once a month*, *once a week*, *several times per week*, *Daily*) for rating.

The macronutrient composition (i.e., percentages of proteins, fats, and carbohydrates per portion) for each participant’s last meal, as well as its total caloric content (kcal), was determined using data from the Food and Nutrient Database (https://fddb.info/db/en/) and the Dutch National Food Consumption Survey 2007–2010 (van Rossum et al., 2011). This procedure involved four steps: 1) If the reported meal did not specify quantities in grams, portion sizes for each food item were estimated using median consumption values from the Dutch survey, taking the participant’s gender into account. 2) These estimated quantities were converted into their respective macronutrient contributions. 3) The grams of each macronutrient were multiplied by its Atwater Factor (Southgate & Durnin, 1970) (4.0 kcal/g for proteins, 9.0 kcal/g for fats, and 4.0 kcal/g for carbohydrates) (Charrondiere et al., 2004), and these values were summed to obtain the total meal energy. 4) Finally, each macronutrient’s caloric contribution was divided by the total meal energy and multiplied by 100 to yield its percentage of total energy. To ensure reliability, two observers independently performed these calculations using this procedure. The interobserver agreement was strong (for proteins: Pearson *r*(96) = .87; carbohydrates: *r*(96) = .86; fat: *r*(96) = .80; caloric content: *r*(96) = .91).

### Statistical analysis

We constructed different 2 × 2 statistical models, incorporating one within-subject factor and one between-subject factor, to assess whether food distractor images in the task induced an attentional blink at Lag 3. Additionally, *t* tests were performed to further analyze significant interactions and to examine similarities in the perceptual and energetic properties of food items, including *memory recall*, *hunger levels*, *explicit food liking*, *consumption frequency*, *perceived caloric content*, and *explicit food craving*. T tests were also used to compare the number of memorized food items across the two food-type conditions. ANCOVAs were performed to evaluate participants’ performance in series with distractors, considering *Lag* (Lag-3 vs. Lag-9) and *food type* (savory food vs. sweet food) as factors. BMI was included as a covariate in all analyses to account for its potential influence on the model outcomes. This adjustment was considered necessary given the broad BMI range observed in all of the experiments, which included several individuals with BMI values outside the normal weight range (< 18 and > 25). We report effect sizes as Cohen’s d for *t* tests and partial eta-squared values for ANOVAs and ANCOVAs. The effect of the covariate was reported only when it reached statistical significance or significantly interacted with other factors.

Finally, we analyzed the impact of the variables *distractor salience, BMI*, and the subjective measurements that we found to be distinctive for *food-type* conditions on the EAB via generalized linear mixed modeling (GLMM) in R (version 4.1.2) and RStudio (http://www.rstudio.com) with the lme4 package (Bates et al., 2015). All of the models we tested included the intercepts of both the participants and the items (the distractor images) as random effects. We first constructed our simplest model, which included the *Lag* and *food type factors* as fixed effects. When the interaction term between these two factors was statistically significant in the ANOVA, we included it in the model. We then included *distractor salience* and *BMI,* one by one, in the model as fixed effects to test their contribution to the EAB effect. Finally, we included the variables encoding the perceptual properties of food images, but only when the *t* test that we conducted revealed differences between the sweet and savory foods. The overall fit from the GLMMs was assessed using *p* values obtained via the likelihood ratio test for each model with the effect against the same model without the effect. Additionally, the Akaike information criterion (AIC) was also used to select the model with the best fit.

## RESULTS

### Memory task performance

We confirmed the effectiveness of the memory task in presetting a mindset. Accordingly, the participants were able to report approximately 6 out of 9 memorized food images correctly at the end of the experiment (*M* = 6.11 ± 1.65). We also observed that savory food images were easier to remember than sweet ones [Sweet: *M* = 5.41 ± 1.39; Savory: *M* = 6.76 ± 1.61; *t*(94) = 4.37; *p* < 0.001; *d* = 0.89]. However, the number of food images seen within the series and reported after completing the RSVP task did not differ significantly [*distractor salience*, sweet: *M* = 3.00 ± 1.83; savory: *M* = 3.82 ± 2.52; *t*(94) = 1.84; *p* = 0.070; *d* = 0.37].

### EAB elicited by sweet and savory foods in the RSVP task

The main effect of food type was not significant [*F*(1,93) = 0.43; *p* = 0.514; η_p_^2^ = 0.005]. Crucially, the target detection accuracy was significantly lower in trials where the distractor image appeared in Lag-3 than in trials where the distractor was placed in Lag-9 [Lag-3: *M* = 90.22 ± 8.78; Lag-9: *M* = 93.61 ± 7.31; *F*(1,93) = 4.09; *p* = 0.046; η_p_^2^ = 0.042]. This effect was modulated by food type [*food type × lag*: *F*(1,93) = 6.82; *p* = 0.011; η_p_^2^ = 0.068], as a larger accuracy difference between Lag-3 and Lag-9 was observed for savory foods [Lag-3: *M* = 88.85 ± 9.59; Lag-9: *M* = 93.89 ± 6.69; *t*(49) = -4.48; *p <* .001; *d* = -0.63; sweet foods: Lag-3: *M* = 91.71 ± 7.64; Lag-9: *M* = 93.32 ± 8.01; *t*(45) = -2.15; *p =* 0.037; *d* = -0.32; see Figure 2].

**Figure 2.**
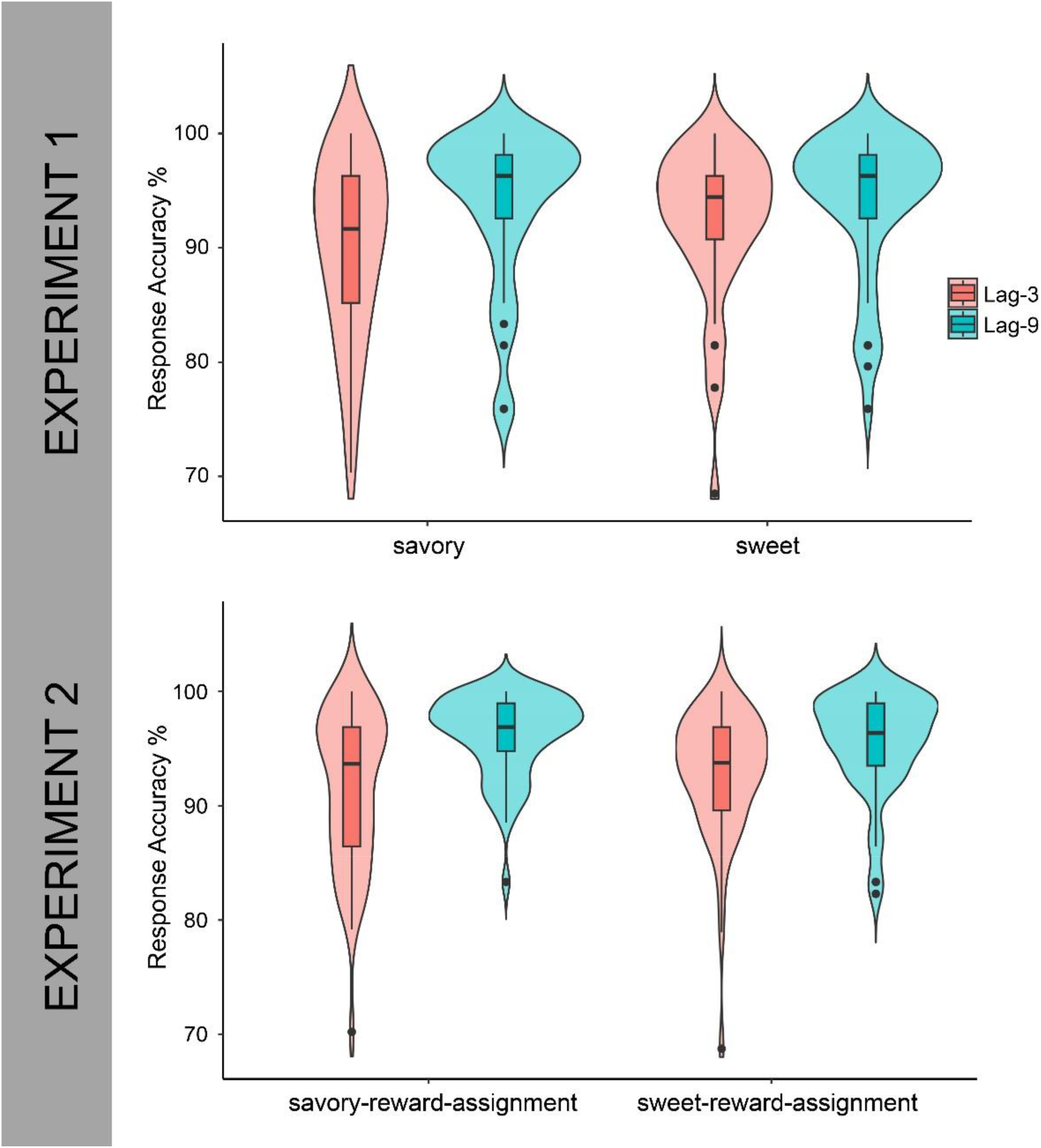
RSVP task performance (Lag 3 vs. Lag 9) in Experiments 1 and 2. In Experiment 1, the results are separated by food-type condition (savory vs. sweet), whereas in Experiment 2, the separation corresponds to the food-reward-assignment condition (savory vs. sweet).

### The EAB and stimuli salience

The EAB effect was included in product‒moment correlation analyses with the measurements possibly associated with stimuli salience in the current task (see methods for details). None of the computed correlations yielded significant results (all *p values > 0.1*). The product‒ moment correlation of EAB with the percentage of macronutrients consumed by participants in their last meal revealed that only the *protein* percentage intake for the savory foods reached a significant negative correlation [savory foods: *r*(48) = -.30; *p* = 0.036; sweet foods: *r*(44) = 0.05; *p =* 0.723]. In other words, we found that EAB increased as the amount of protein ingested in the last meal decreased (see Figure 3). In contrast, neither carbohydrate nor fat percentages nor the total amount of calories consumed were found to be correlated with the ABias^1^. All other subjective ratings were not correlated with EAB (all *p values* > 0.20).

**Figure 3.**
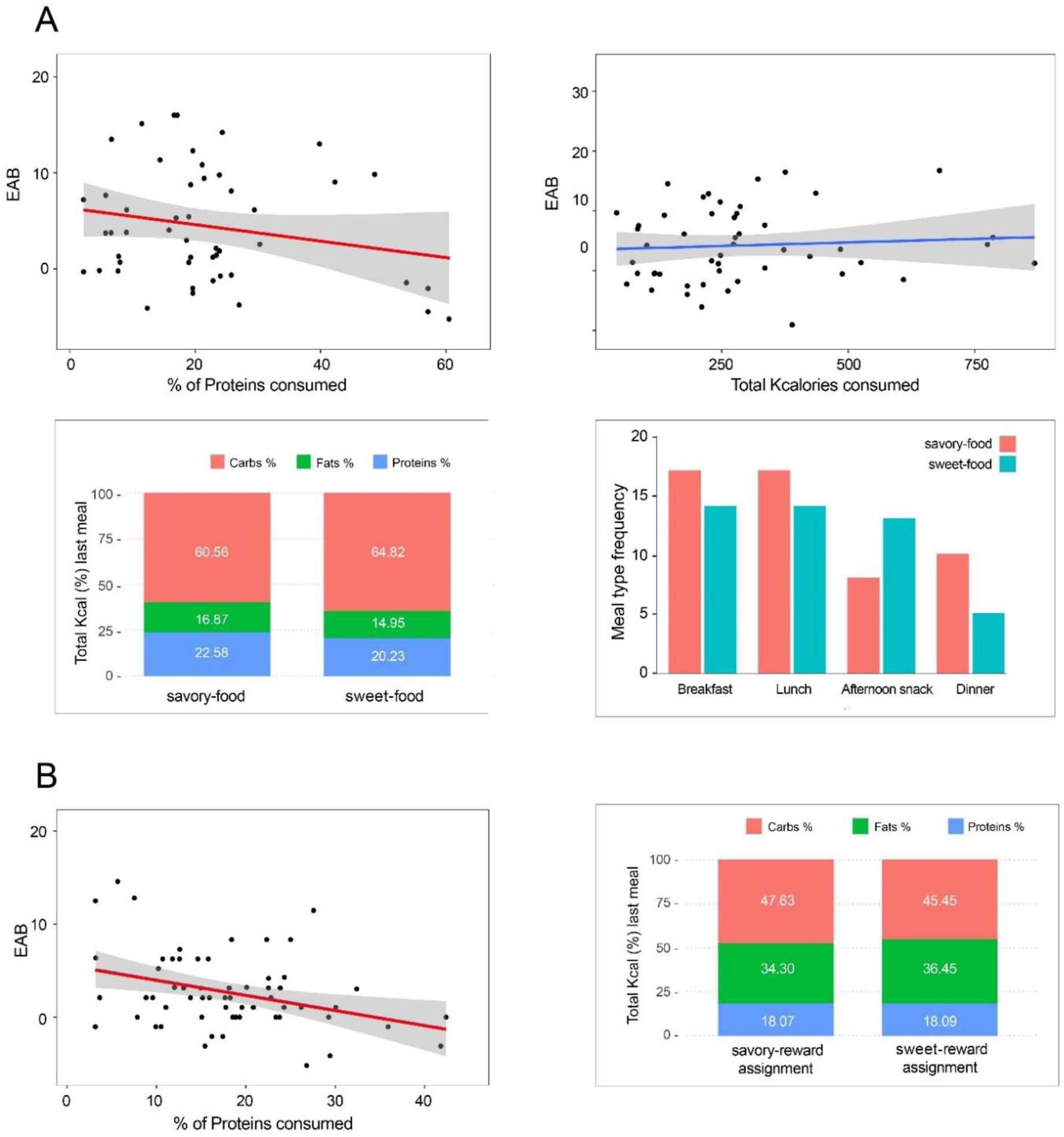
Correlations, macronutrient distributions, and last meal intake in Experiments 1 and 2. A. In Experiment 1 (savory-food condition), a significant negative correlation [*r*(50) = -.33, *p* = .016] was found between the percentage of protein consumed in the last meal and the EAB (Lag-9 minus Lag-3), indicating that lower protein intake before the experiment was associated with a greater EAB. No significant differences were detected in the distribution of macronutrients (bottom left) or in the frequency of last meal types (breakfast, lunch, afternoon snack, and dinner) between participants assigned to the savory-food conditions and those assigned to the sweet-food conditions [χ^2^(3) = 3.08, *p* = .38] (bottom right). **B.** (left) In Experiment 2 (sweet-reward-assignment condition), a similar negative correlation emerged between protein percentage and EAB [*r*(60) = -0.35; *p =* 0.006]. No significant differences were found in the macronutrient distributions in the last meal between the savory and sweet conditions (see Table 2S).

**Table 2.** Summary of subjective ratings of images’ perceptual food properties for Experiments 1 and 2.

| EXPERIMENT-1 | sweet-food | savory-food | sweet vs. savory |  |
| --- | --- | --- | --- | --- |
|  | Mean (SD) | Mean (SD) | t-value (df) | p-value |
| Hunger feeling | 4.23 (2.7) | 3.52 (2.9) | 1.23 (94) | 0.22 |
| Explicit liking | 6.68 (1.5) | 6.96 (1.3) | 0.97 (94) | 0.33 |
| Consumption frequency | **5.12 (0.6)** | **4.54 (0.6)** | **4.50 (94)** | **<0.001** |
| Perceived calories | 8.07 (1.0) | 7.70 (1.05) | 1.78 (94) | 0.07 |
| Explicit craving | 3.95 (2.1) | 3.67 (2.4) | 0.74 (94) | 0.54 |

| EXPERIMENT-2 | sweet-reward-assig. | savory-reward-assig. | sweet vs. savory |  |
| --- | --- | --- | --- | --- |
|  | Mean (SD) | Mean (SD) | t-value (df) | p-value |
| Hunger feeling | 7.14 (1.8) | 7.32 (1.4) | -0.59 (115) | 0.56 |
| Explicit liking sweet | 7.08 (2.1) | 7.08 (2.1) | 0.02 (115) | 0.98 |
| Explicit liking savory | 6.28 (1.7) | 6.69 (1.9) | -1.25 (115) | 0.22 |
| Consump. freq. sweet | 4.98 (0.7) | 4.83 (0.9) | 1.02 (115) | 0.31 |
| Consump. freq. savory | **4.63 (0.8)** | **4.26 (0.8)** | **2.37 (115)** | **0.019** |
| Perc. calories sweet | 8.50 (1.2) | 8.36 (1.2) | 0.67 (115) | 0.51 |
| Perc. calories savory | 7.47 (1.2) | 7.30 (1.2) | 0.76 (115) | 0.45 |
| Explicit craving sweet | 5.45 (2.6) | 5.32 (2.4) | 0.28 (115) | 0.78 |
| Explicit craving savory | 5.48 (1.9) | 5.80 (2.2) | -0.84 (115) | 0.40 |

**Note.** Hunger ratings reflect each participant's mean hunger level before and after the experiment.

Finally, among all of the ratings (see Table 2), only consumption frequency significantly differed, indicating that participants consumed sweet foods more often than savory foods did [*t*(94) = -4.49; *p <* 0.001; *d* = -0.92].

We subsequently analyzed whether consumption frequency or distractor salience could explain the EAB effect in our RSVP task. A model including *Lag*, *food type*, and *consumption frequency* as fixed effects, together with the two-way interactions between *Lag* and the remaining predictors, was compared against the simplest model (fixed factors: *Lag*, *food type*, and lag × *food type*). A comparison of these two models revealed that the inclusion of *consumption frequency* in the model did not improve its fit [χ^2^(2) = 1.24; *p* = 0.539]. However, we did observe an improvement in the model fit when *distractor salience* was included [χ^2^(2) = 31.51; *p* < 0.001]. The model that better explained the EAB (marginal *R^2^* = 0.04; conditional *R^2^* = 0.23), however, included the fixed factors *Lag, food type, distractor salience,* and *BMI* (Table 3), as well as the two-way interactions between *Lag* and all of the other predictors described here (Table 4)^2^.

**Table 3.** Akaike information criterion (AIC) scores of the different models tested for Experiments 1 and 2.

**EXPERIMENT 1**
| *Model formula* | *AICc* | *ΔAICc* | *W<sub>i</sub>* |
| --- | --- | --- | --- |
| Hits ~ *Lag* * (*food-type* + BMI + *distractors-awareness*) + (1\|ID) + (1\|Item) | 5350.1 | 0.00 | 0.88 |
| Hits ~ *Lag* * (*food-type* + *distractors-awareness*) + (1\|ID) + (1\|Item) | 5354.1 | 3.98 | 0.12 |
| Hits ~ *Lag* * *food-type* + (1\|ID) + (1\|Item) | 5381.6 | 31.48 | 0.00 |
| Hits ~ *Lag* + *food-type* + (1\|ID) + (1\|Item) | 5387 | 36.87 | 0.00 |

**EXPERIMENT 2**
| *Model formula* | *AICc* | *ΔAICc* | *W<sub>i</sub>* |
| --- | --- | --- | --- |
| Hits ~ *Lag* * (*food-reward assignment* + *distractors-awareness*) + (1\|ID) | 9683.04 | 0.00 | 1 |
| Hits ~ *Lag* * *food-reward assignment* + (1\|ID) | 9697.46 | 14.42 | 0 |
| Hits ~ *Lag* * (*food-reward assignment* + *Consumption Frequency*) + (1\|ID) | 9700.36 | 17.31 | 0 |
| Hits ~ *Lag* * (*food-reward assignment* + BMI) + (1\|ID) | 9701.27 | 18.23 | 0 |

**Note.** The smaller the AIC value is, the better the model fit. The **delta AICc** represents the difference in the AIC score between the best model and the model compared, whereas the **W<sub>i</sub> AICc** weight indicates the proportion of predictive power provided by the full set of models contained in the model tested. Convergence was reached in all of the models.

**Table 4.** Logistic linear mixed model estimated using ML for Experiments 1 and 2.

**EXPERIMENT 1**
| *Predictors* | *Odds Ratios* | *CI* | *Estimate* | *p-value* |
| --- | --- | --- | --- | --- |
| (Intercept) | 12.23 | 8.90 – 16.81 | 15.44 | <0.001 |
| **Lag [Lag-9]** | **1.82** | **1.48 – 2.24** | **5.65** | **<0.001** |
| Food-type [Sweet] | 1.23 | 0.81 – 1.86 | 0.97 | 0.331 |
| BMI | 0.98 | 0.80 – 1.21 | -0.15 | 0.880 |
| **Distractors-awareness** | **0.64** | **0.52 – 0.79** | **-4.22** | **<0.001** |

<sup>2</sup> We also included in the basic model, one by one, the factors corresponding to the distinctive perceptual properties of foods, and the number of memorized images recalled at the end of the experiment. None of those factors were found to improve the model fit when they were incorporated in the model.

|  |  |  |  |  |
| --- | --- | --- | --- | --- |
| Lag [Lag-9] * food-type [Sweet] | **0.71** | **0.52 – 0.95** | **-2.27** | **0.023** |
| Lag [Lag-9] * BMI | **0.82** | **0.70 – 0.95** | **-2.66** | **0.008** |
| Lag [Lag-9] * distractors-awareness | **1.44** | **1.24 – 1.67** | **4.85** | **<0.001** |

EXPERIMENT 2
| *Predictors* | *Odds Ratios* | *CI* | *Estimate* | *p-value* |
| --- | --- | --- | --- | --- |
| **(Intercept)** | 21.32 | 13.13 – 34.62 | 12.37 | <0.001 |
| Lag [Lag-9] | 1.37 | 0.99 – 1.89 | 1.91 | 0.056 |
| Food-type [Sweet] | 1.29 | 0.91 – 1.83 | 1.43 | 0.154 |
| **Distractors-awareness** | **0.91** | **0.84 – 0.99** | **-2.15** | **0.032** |
| **Lag [Lag-9] \* food-type [Sweet]** | **0.66** | **0.53 – 0.84** | **-3.45** | **0.001** |
| **Lag [Lag-9] \* distractors-awareness** | **1.13** | **1.07 – 1.19** | **4.19** | **<0.001** |

**Note.** All *p* values were computed via the Wald approximation. The values depicted in bold indicate statistically significant effects of the predictors.

## DISCUSSION

In Experiment 1, we aimed to determine whether ABias toward high-caloric food serve as an all-purpose cognitive function to facilitate the acquisition of highly energetic supplies or could otherwise be interpreted as facilitating the attainment of specific food types. To that end, we effectively preset the mindset of the participants, with images of sweet or savory foods increasing in that way in salience. Then, we tested whether they elicited a similar EAB when used as distractors in an RSVP task. Our results revealed superior ABias toward foods representative of an elevated protein content. Savory foods were more salient than sweet foods were, as they were much more efficient in capturing attention in a task in which all food images presented were completely irrelevant to solve the task. The attentional attraction toward proteins was further supported by the larger protein-related ABias observed when the percentage of proteins consumed in the last meal was low. These results indirectly highlight the importance of protein consumption in preserving metabolic homeostasis in omnivores (Jensen et al., 2012; Simpson et al., 2004) and suggest that the brain may prioritize redirecting attentional resources to facilitate the acquisition of essential amino acids when an opportunity arises.

After attempting to elucidate the origin of the observed EAB effect in Experiment 1, we deemed it necessary to replicate our main results, particularly those encountered for high-energy foods rich in proteins.

## EXPERIMENT 2

In the previous experiment, we elucidated that to induce EAB using food images as distractors in an RSVP task, these stimuli must attain sufficient salience to capture participants’ attention. Although we successfully manipulated stimuli salience by instructing participants to memorize food stimuli subsequently used as distractors in the RSVP task, other similar variants of the task have proven effective in eliciting EAB for task-irrelevant stimuli. For example, images of an angry face presented at Lag-2 within a series of images (well within the temporal window where the EAB occurs) disrupted task performance (Gutiérrez-Cobo et al., 2019). In that experiment, participants were explicitly informed that emotional stimuli presented in the series of images indicated a reward contingent on a correct response in those trials. In a subsequent experiment, even when instructions were modified to focus on identifying the gender of the face rather than directly attending to the emotion for reward determination, similar EABs were consistently observed. Thus, with this other variant of the RSVP task, the authors were able to increase the salience of the facial images while maintaining them in a state of irrelevancy for completing the task. Following a similar stimulus salience manipulation, we conducted a second experiment to further investigate the influence of the macronutrient compositions of food cues on the EAB concerning high-caloric foods. Thus, in Experiment 2, we removed the memory task and explicitly instructed participants that the presence of some food images within the series indicated a contingent reward trial.

## METHODS

### Experiment 2

#### Participants

A new group of 123 students from the Faculty of Psychology at the University of Barcelona, none of whom had participated in the previous experiments, took part in the present laboratory-based experiment. All of the participants provided informed consent upon enrollment. We excluded data from 6 participants because their performance in conditions without the distractor was less than 2.5 S.D. from the mean. The final sample comprised 117 participants, who were randomly and automatically assigned to the savory-reward (57) or sweet-reward (60) food-type condition upon entering the experiment. The two samples were similar in terms of age, BMI (*p* values > 0.2), and gender composition [χ^2^(1) = 2.74; *p* = 0.098] (Table 1). The participants were compensated with course credits and received a monetary bonus on the basis of their performance.

### Stimuli and procedure

All of the stimuli were identical to those used in the previous experiments, with the exception that the food stimuli were limited to 4 images for each food category from the same pool of 18 food images used in the previous experiments (sweet foods: vanilla ice cream cone, Oreo’s cookies, chocolate popsicle with nuts, and pralines; savory foods: burger with bacon, cheese platter, salami sausage, and meat pie). We conducted a new set of statistical analyses to ensure that the energetic properties of the two food sets were comparable, except for the sugar and protein contents (see the Supplementary Material).

The procedure differed from the previous experiments in that the participants were directly engaged in the RSVP task without previously undertaking any other task. The participants were required to attend to the experiment following a fasting period of at least 5 hours (*M* = 11.6; SD = 3.7). They were instructed, as in the previous experiment, to report in each trial— at the end of the sequence of images—the appearance or absence of the target stimuli, using the mouse buttons. Additionally, the eight food images were presented to them in the center of the screen, which was arranged randomly in two rows of four images. The participants were informed that they could earn points for correct responses in trials where some of those food images were presented and that they would have to determine which were the rewarded ones. However, they were also told that to maximize their monetary gains, they should focus on performing the task as accurately as possible.

The participants were subsequently informed that each reward trial would result in a 50-point gain for a correct response and a 50-point loss for an error. On trials where the food image was not rewarded, i.e., images belonging to the other food category, either sweet or savory foods, they were informed that they would not gain or lose points for providing a correct or incorrect response. Finally, the participants were told that the number of points accumulated at the end of the experiment would determine their monetary bonus, with a maximum set at 10€. For half of the participants, sweet foods were considered valued distractors, while savory foods were considered nonvalued distractors, and vice versa. At the beginning of the experiment, each participant was randomly and automatically assigned to a condition that was not disclosed to them. The experiment was programmed in Python (version 3.12.7), using the Pyglet library for stimulus presentation and interaction (Holkner & Pyglet-developers, 2021), and presented on a 24” LCD monitor with the refresh rate set at 120 Hz. The participants comfortably sat in front of a computer screen, with the stimuli presented subtending at approximately 5.7° degrees of the visual angle.

The RSVP task consisted of 8 blocks, each containing 72 trials (total of 576), distributed among 24 valued, 24 nonvalued, and 24 neutral trials presented in random order. The arrangement of images within each trial and the general procedure throughout the experiment were identical to those used in Experiment 1. This time, however, feedback was provided contingent on each response, lasting for a duration of 833 ms. The information displayed on the screen varied on the basis of both the condition and the accuracy of the participant’s response in that trial. Thus, for correct responses in valued trials, “CORRECT: YOU WIN 50 POINTS!!” is displayed in green, whereas for incorrect responses, “ERROR: YOU LOSE 50 POINTS” is displayed in red. In neutral and nonvalued trials, the information displayed was limited to "CORRECT" or "INCORRECT," in black, depending on the accuracy of the response. Short breaks were incorporated at the end of each block, which participants ended by pressing the space bar. Finally, after the experiment, participants were shown their monetary bonus, calculated as 5.5 + (points – 3,780)/1,890.

The same subjective measures were collected as in the previous experiments; however, the memory task-related questions were replaced by the following items: i) list all of the food images you have seen in the previous task, ii) list all of the food images identified as yielding the reward (*valued items*), and iii) list all of the food images identified as not yielding the reward (*nonvalued items*). Finally, in addition to reporting the foods consumed in their most recent meal, the participants were also asked to report what they had eaten for dinner the previous day, given that the experiment was conducted between 12 p.m. and 4 p.m. The entire experiment lasted approximately 40 min.

After the experiment, two observers independently calculated the macronutrient composition of the whole foods of the participants’ last meal and last dinner using the procedure described in Experiment 1. The interobserver agreement was again strong in all of the cases [last meal, proteins: Pearson *r*(117) = .81; carbohydrates: *r*(117) = .83; fat: *r*(117) = .80; caloric content: *r*(117) = .89; dinner, proteins: *r*(117) = .82; carbohydrates: *r*(117) = .83; fat: *r*(117) = .83; caloric content: *r*(117) = .91].

## RESULTS

### RSVP task performance

The participants were 96.0% accurate in sequences without the distractor (neutral condition), indicating correct task implementation and performance. They earned an average of 8.1€, with no significant differences between groups [*t*(115) = 1.54; *p* = 0.127; *d* = 0.28].

By the end of the experiment, the participants reported seeing an average of approximately five of the eight food images in the RSVP sequences (M = 5.0 ± 2.08). Notably, they identified food-valued items significantly more often than nonvalued items [*F*(1,115) = 10.30; *p* = 0.002; η_p_^2^ = 0.082]. Neither the main effect of food-reward assignment [*F*(1,115) = 2.58; *p* = 0.111; η_p_^2^ = 0.022] nor the interaction between food value and food-reward assignment [*F*(1,115) = 0.36; *p* = 0.551; η_p_^2^ = 0.003] reached significance. Collectively, these findings support the successful implementation of this version of the RSVP task, indicating no a priori differences between the two groups of participants while highlighting an advantage for remembering reward-related food– items.

### EAB effect elicited by sweet-value and savory-value foods in the RSVP task

Next, we evaluated whether the food distractor images captured attention. The ANCOVA results revealed a significant lag × *food-reward assignment* interaction [*F*(1,114) = 8.27; *p* = 0.005; η_p_^2^ = 0.068]. Neither the main effect of Lag [*F*(1,114) = 2.01; *p* = 0.159; η_p_^2^ = 0.017], *food value* [*F*(1,114) = 0.01; *p* = 0.94; η_p_^2^ < 0.001], *food-reward assignment* [*F*(1,114) = 0.71; *p* = 0.400; η_p_^2^ = 0.006], nor any other interaction term reached statistical significance (all *p* > 0.4) (Figure 2). Follow-up analyses revealed that the attentional capture of the food distractor’s image was observed in both *food-reward-assignment* conditions [sweet-reward- assignment conditions, Lag-3: *M* = 92.87 ± 5.50; Lag-9: *M* = 95.57 ± 4.40; *t*(59) = -5.11; *p* < 0.001; *d* = -0.66; savory-reward-assignment, Lag-3: *M* = 90.62 ± 8.08; Lag-9: *M* = 96.23 ± 3.55; *t*(56) = -6.38; *p* < 0.001; *d* = -0.85], although as reflected in the EAB, i.e., the difference between Lag-3 and Lag-9, such an effect was greater for participants in the savory-reward- assignment [*t*(115) = -2.87; *p* = 0.005; *d* = -0.53]. Overall, these findings suggest that the reward manipulation implemented in Experiment 2 effectively increased stimulus salience— thus capturing attention and reducing task performance—while also replicating the stronger attentional capture for savory foods than for sweet foods observed in Experiment 1.

### The EAB and stimuli salience

We calculated ABias from the current dataset and examined it using Pearson correlation analyses to determine whether the amount of the different macronutrients consumed in the last meal influenced the EAB, as suggested by Experiment 1. Among the three macronutrients, only *protein* intake showed a significant negative correlation in participants with sweet food assigned as value task items [*r*(60) = -0.35; *p =* 0.006], indicating that a higher EAB was associated with lower protein consumption (see Figure 3). No significant correlations were observed in the savory-reward assignment (all *p* values > 0.1; Figure S2). Similarly, the same product‒moment correlation analyses of macronutrient percentages consumed at dinner on the previous day did not yield any significant results.

Next, food image properties, evaluated by all of the participants at the end of the experiment, were compared across conditions using separate ANOVAs with reward assignment (savory- reward vs. sweet-reward) as the between-subjects factor and food type (savory vs. sweet) as the within-subjects factor.

For explicit liking, the participants rated the sweet food images (*M* = 7.08 ± 2.06) higher than the savory foods did (*M* = 6.48 ± 1.78) [*F*(1,115) = 6.92; *p* = 0.010; η_p_^2^ = 0.057], regardless of food-reward assignment [*F*(1,115) = 0.53; *p* = 0.470; η_p_^2^ = 0.005; *food-reward assignment × food type*: *F*(1,115) = 0.85; *p* = 0.357; η_p_^2^ = 0.007]. The craving scores did not differ significantly across conditions (all *p* values > 0.3). However, the participants perceived sweet foods as more caloric [*F*(1,115) = 59.26; *p* < 0.001; η_p_^2^ = 0.340] and reported consuming them more frequently than savory foods did [*F*(1,115) = 20.11; *p* < 0.001; η_p_^2^ = 0.149]. Furthermore, participants assigned to the sweet-reward condition reported consuming all tested foods more often than those in the savory-reward condition did [*F*(1,115) = 5.34; *p* = 0.023; η_p_^2^ = 0.044]. No other main effect or interaction terms reached statistical significance (all *p* values > 0.3).

Following the analytical procedure of Experiment 1, we subsequently estimated whether distractor salience (food items reported to have been seen during the RSVP task), explicit food liking, perceived calories, and food consumption could explain the EAB effect in this version of the RSVP task. We conducted generalized linear mixed models (GLMMs) to assess the relevance of *those variables* to our results. A model including *Lag*, BMI, and *consumption frequency* as fixed effects, together with their interactions between *Lag* and the remaining predictors, was compared against the simplest model (fixed factors: *Lag*, food-reward assignment, and *Lag* x food-reward assignment). All of the models we tested included the intercept for participants as a random effect.^3^ A comparison of these two models revealed that the inclusion of *consumption frequency* in the model did not improve its fit [χ^2^(2) = 1.11; *p* = 0.575] or BMI [χ^2^(2) = 0.192; *p* = 0.909]. We again observed an improvement in the model fit when *distractor salience* was included in the model [χ^2^(2) = 18.42; *p* < 0.001], the same observation as in Experiment 1. The model that better explained the EAB (marginal *R^2^* = 0.04; conditional *R^2^* = 0.21) included the fixed factors *Lag, food-reward assignment, and distractor salience* (Table 3), as well as the two-way interactions between *Lag* and all of the other predictors described here (see Table 4 for a summary of the results of the tested models).

## DISCUSSION

The results of Experiment 2 highlight the importance of stimulus salience and the protein content of food images in eliciting an EAB during the RSVP task. In this experiment, the participants’ mindset was preset by simply informing them that certain food images indicated a contingent reward for accurate responses; however, they were advised to focus on target detection to maximize monetary gains. Nonetheless, participants’ task performance was disrupted when food image distractors appeared close before the target, i.e., in the Lag-3 condition, but not when it was presented outside the time windows where EAB is known to occur. Moreover, when savory (protein-rich) foods were the valued images, the attentional disruption was greater than when sweet foods were valued, replicating the findings of Experiment 1 and highlighting again the role of the macronutrient composition of food cues in eliciting an EAB. Additional evidence for the influence of protein in capturing attention emerged from the correlation between protein intake in the last meal and the extent of attentional disruption, with lower protein consumption associated with stronger EAB for the participants for which sweet food indicated rewarding trials; this relationship did not emerge when savory (protein-rich) foods were already valued.

## GENERAL DISCUSSION

In the present study, we aimed to investigate the deterministic role of mindset in modulating the salience of food cues to capture attention and to explore a possible physiological function underlying such attentional modulation. Specifically, we sought to determine whether an active mindset was sufficient and necessary for inducing attentional capture. Additionally, by using different sets of high-calorie foods differentiated by their macronutrient composition, we examined whether food-related EAB followed solely an energetic supply function or could alternatively be considered to facilitate the selection of foods with specific macronutrients to preserve nutritional homeostasis.

We found in Experiment 1 that the induction of an active mindset in the RSVP task was effective in modulating the attentional capture of food images (see also supporting experiments in the Supplementary Material). In Experiment 2, we omitted the memory task; instead, we used food images in the RSVP task to merely signal reward availability. This was done to increase the salience of those food images and thereby evoke an EAB. We found that attributing an explicit value to the otherwise task-irrelevant food images in the RSVP task was sufficient to increase their salience and to capture participants’ attention, even though participants were explicitly instructed to avoid attending to such informative items to maximize their task accuracy.

On the one hand, certain food features, such as appeal, have been shown to influence food choices and consumption (Birch, 1999; Marty et al., 2018). Highly palatable foods can “rewire” mesolimbic dopamine neurons, leading to food-seeking behaviors even after limited exposure (S. Liu et al., 2016). Hence, cumulative food experience may be at least partially responsible for modulating ABias toward food (Anderson et al., 2021). This idea is supported by the phenomenon of value-modulated attentional captures (Anderson et al., 2011), as evidenced across multiple experiments, which indicates that learning about the reward associated with stimuli modulates attention to those stimuli (for a review, see (Le Pelley et al., 2016; Pearson et al., 2022)). At the experimental level, it has recurrently been shown that attention is rapidly and automatically directed toward or taken away from stimuli that have been repeatedly paired with rewarding or aversive outcomes, respectively, throughout individuals’ learning experiences. This value-related attentional phenomenon has been formalized as an integral component of attentional control known as *selection history* (Anderson et al., 2021), which complements the well-known dichotomous mechanism (top-down and bottom-up) of attentional control (Corbetta & Shulman, 2002). Hence, selection history appears well positioned to elucidate why palatable food items can capture attention more effectively than less common food items when they are part of a mindset (Ballestero-Arnau et al., 2021).

On the other hand, a physiological benefit of learning attention toward food, especially high-caloric food, may correspond to the need to guarantee an optimal energy supply. This justifies the underlying research motivation for employing high-caloric foods when examining different factors related to food ABias, such as hunger and obesity. However, cumulative evidence from the last two decades seems to indicate the existence of physiological self-regulatory mechanisms for attaining a sufficient level of salt and protein intake (Seeley & Berthoud, 2000). No mechanism has been described yet to detect specific deficits in carbohydrates and fat, perhaps because most carbohydrates and lipids can be synthesized internally. Nutritional state-dependent self-regulation of protein intake has been shown in other animals (Chaumontet et al., 2018; Murphy et al., 2018), with low protein consumption causing an increase in protein intake in subsequent meals (Raubenheimer & Simpson, 2019; Simpson & Raubenheimer, 2014a), a hypothesis that has received substantial support from neuroimaging studies (Gosby et al., 2011; Martinez-Cordero et al., 2012; Raubenheimer & Simpson, 2019; Simpson & Raubenheimer, 2005, 2014b).

Currently, the overwhelming existence of ultra-processed foods with high sugar contents allows easy access to a rich source of energy in the form of rapidly absorbed carbohydrates (Kellett et al., 2008; Roden & Bernroider, 2003). In contrast, proteins are assumed to play a primary role in nutrition (Raubenheimer & Simpson, 2019; Simpson & Raubenheimer, 2005) since essential amino acids cannot be synthesized internally and are crucial for health maintenance, growth, reproduction and immunity (Wu, 2009). In this context, the protein leverage hypothesis (PLH) postulates that humans prioritize the ingestion of proteins over carbohydrates and fats (Martinez-Cordero et al., 2012). Interestingly, PLH predicts the overeating of nutrients with low protein concentrations—although rich in fats and carbohydrates—due to the need to reach an optimal consumption of proteins (Griffioen-Roose et al., 2014; Raubenheimer & Simpson, 2019), a phenomenon recently associated with the rise and prevalence of obesity (Gosby et al., 2014; Hall, 2019; Martínez Steele et al., 2018). However, although the PLH has received relevant support from randomized controlled trials (Campbell et al., 2016; Gosby et al., 2011; Hall et al., 2019; Kohanmoo et al., 2020) and neuroimaging studies (Griffioen-Roose et al., 2014; Leidy et al., 2011), supporting evidence at the cognitive level of specific attentional prioritized protein stimuli is still lacking.

In the present study, we found that individuals displayed increased ABias toward protein-rich foods and that the magnitude of the ABias was negatively correlated with the percentage of proteins ingested in their preceding meal. To our knowledge, this is the first evidence that ABias toward food might underlie a physiological response associated with nutritional homeostasis. Nevertheless, the novelty of our findings may simply stem from the oversight of previous studies, which did not examine participants’ dietary intake before the experiment and omitted controlling for the macronutrient composition of the food cues used. Food and food-related stimuli are considered inherently salient, with ABias toward food cues reflecting the significant impact they have on the perceptual system, as they are typically more easily noticed and preferentially selected. Accordingly, the automaticity of such ABias may be directly linked to the effective salience of food-related stimuli in terms of predictive value, created as a result of accumulative (valued) life experience with food and its nutritive consequences. In support of this notion, it has been shown that exposure to stimuli representing high-protein content, after a low-protein state is provoked, induces accentuated activation of reward-related brain regions (Griffioen-Roose et al., 2014; Simpson et al., 2003; Simpson & Raubenheimer, 2005). The idea that protein-related cues may be regarded as highly salient stimuli in conditions of low protein availability bears a close resemblance to the concept of alliesthesia (Cabanac, 1971). With respect to food perceptual properties, alliesthesia posits that the rewarding value of a food-related stimulus is influenced by the internal state of the organism, such as hunger or satiety (Cabanac & Fantino, 1977; Jiang et al., 2008; Plailly et al., 2011). Similarly, the state of hunger has been directly associated with the motivational significance of stimuli (Hardman et al., 2021) and is suggested to affect cognitive processes such as perception and attention (Al-Shawaf, 2016).

It seems that we are currently facing a paradox that what may constitute an essential physiological phenomenon underlying human evolution, i.e., the regulation of protein intake (Richards & Schmitz, 2008; Richards & Trinkaus, 2009), may have turned against us. The low protein content found in almost all ultra-processed foods appears to be strongly associated with the recent emergence of the obesity pandemic (Hall, 2019; Hendrikse et al., 2015; Simpson & Raubenheimer, 2005, 2014b; Swinburn et al., 2011). Alternatively, at a speculative level, the observed enhancement of the ABias toward protein-related foods that in this study might simply be indicative of a particular sensitivity to meat consumption in our test population, which may be aligned with current cultural movements in support of animal rights and against climate change (Fiala, 2008; Parker et al., 2018). Overall, given the current findings and to investigate the possible causal relationship between habitual low protein intake in the diet and increased ABias toward high-protein foods, future studies should explore the feasibility of manipulating participants’ diet (or last meal ingestion) to replicate and validate the present findings.

## SUPPLEMENTARY MATERIAL METHODS

### Stimuli properties

#### Experiment 1

We ensured that the *spatial frequency, brightness, image size, familiarity, valence,* and *arousal* were similar between the two sets of images. The only significant difference we found was for *arousal*, which was greater when comparing food images—all items together—with those from the nonfood category [food vs. nonfood: *t*(35) = 6.54; *p* < 0.001; *d* = 2.15] and *valence*, for which a marginally significant difference was found [*t*(35) = 1.93; *p* = 0.062; *d* = 0.64]. These greater values for *arousal* and *valence* in food images were assumed to be intrinsic defining factors of the food category. Furthermore, and as expected, images of sweet foods had a significantly greater amount of added sugar [sweet vs. savory: *t*(16) = 6.48; *p* < 0.001; *d* = 4.40], as well as carbohydrate [*t*(16) = 3.15; *p* = 0.006; *d* = 1.60], than did those of savory foods, whereas savory foods had more proteins [*t*(16) = 4.13; *p* = 0.002; *d* = 1.9]. The fat content (*p* = 0.79) and caloric density (kcal/100 g; *p* = 0.15) were similar in the two food categories, whereas the amount of salt was slightly greater for savory foods [savory: 0.60 g/100 g; sweet: 0.13 g/100 g; *t*(16) = 2.35; *p* = 0.038; *d* = 1.40].

#### Experiment 2

The results of the Mann‒Whitney U tests revealed that the selected sweet food images, as expected, had significantly higher sugar (*p* = 0.029) and carbohydrate (*p* = 0.029) contents than did the savory food images, whereas the savory food images contained significantly more protein (*p* = 0.029). No significant differences were observed in fat content (*p* = 1), caloric density (kcal/100 g; *p* = 0.886), or salt content (*p* = 0.486).

## RESULTS

### RSVP task—baseline—performance

#### Experiment 1

We verified the correct implementation of the RSVP task in its online version by analyzing participant performance in sequences without distractors. In each of the four experiments, separate analyses of variance (ANOVAs) were conducted with the within-subject factor *task accuracy* (target vs. nontarget) and the between-subject factor *food type* (savory vs. sweet).

In Experiment 1, the results showed that participants exhibited accurate overall task performance, with an accuracy of ∼95% for the question of whether the target was or was not presented within the sequence. The participants’ task performance was significantly better in detecting the absence of the target rather than its presence [target accuracy: *M* = 94.59 ± 5.75; nontarget accuracy: *M* = 96.37 ± 3.18; *F*(1,94) = 6.98; *p* = 0.010; η_p_^2^ = 0.069].

However, performance was similar for the two *food-type* groups [*F*(1,94) = 0.31; *p* = 0.580; η_p_^2^ = 0.003; *food-type* × *target-detection*: *F*(1,94) = 0.49; *p* = 0.487; η_p_^2^ = 0.005]. Overall, these results indicated that the task worked as expected without any a priori task differences between sweet and savory foods.

#### Experiment 2

In Experiment 2, the neutral condition showed no significant difference in RSVP task performance between the target-present and target-absent trials [target accuracy: M = 95.14 ± 4.36; nontarget accuracy: M = 96.95 ± 2.35; *F*(1,114) = 0.18, *p* = 0.675, η _p_^2^ = 0.002]. However, a marginally significant difference was found between participants assigned to sweet-valued versus savory-valued food-reward-assignment conditions [*F*(1,114) = 3.61; *p* = 0.060; η_p_^2^ = 0.031], as well as for the *target-detection x food-reward-assignment* interaction [*F*(1,114) = 3.86; *p* = 0.052; η_p_^2^ = 0.033]. Overall, these results suggest that participants in the sweet-value assignment condition were more accurate at detecting the absence of the target than in detecting its presence in trials without the food distractor.

### Food macronutrient distribution and the EAB

The product–moment (bivariate Pearson) correlations are presented as color heatmaps for Experiments 1 and 2, separately for the food-type and food-reward–assignment conditions (Figure S2). Notably, in both experiments, a significant correlation emerged only between Proteins and Attentional Bias, indexed by the EAB effect. Notably, in Experiment 1, this correlation was observed among participants assigned to memorize high-protein foods, whereas in Experiment 2, in which both savory and sweet food were used as distractor images for all participants, a significant correlation was observed only in the group for whom sweet foods were designated as valued items.

Finally, to control for potential confounders, we compared the macronutrient composition and total caloric intake of the last meal between the two groups of participants (savory- reward–assignment vs. sweet-reward–assignment). No significant differences emerged in any of the direct comparisons in either Experiments 1 or 2 (see Table S2).

## ADDITIONAL EXPERIMENTS

The observed EAB effect in Experiment 1 was intricately linked to the presence of distractor food images, proving effective in capturing participants’ attention. These food images were preloaded into the participants’ minds and actively maintained in memory to complete a subsequent memory recall requirement. Accordingly, the need for an active mindset to elicit an attentional blink toward preloaded stimuli in the mindset was further examined in a subsequent experiment. Thus, the importance of an active mindset for enhancing the salience of food distractor images was further investigated in two follow-up experiments, in which food mindsets were deactivated by resolving the memory recall task before the subsequent RSVP task (Experiment 3) or by presenting a different set of food images rather than the memorized ones in the RSVP task (Experiment 4).

## EXPERIMENT 3

In Experiment 1, we effectively primed the participants’ mindset to increase the salience of food images in the subsequent RSVP task, aiming to capture their attention. Although the participants were aware in advance that these food images were task-irrelevant, they were still important for achieving the main objective of the study, i.e., the recall of memorized food items at the end of the experiment. To investigate whether an active mindset was essential for eliciting an EAB effect in the RSVP task or, in other words, to examine whether eliminating the need to maintain a memory trace active for these items during the experiment diminished the salience of the food distractor images, we conducted a second experiment.

Experiment 2 replicated the conditions of Experiment 1, with the sole exception that participants were instructed to report the memorized items before engaging in the subsequent RSVP task. Consequently, the food images preloaded into the participants’ mindset became goal-irrelevant during the RSVP task, as memory recall was already complete before the task. We hypothesize that if an active mindset was crucial for observing an EAB toward the stimuli primed in the participants’ mindset, we would not observe an EAB when these images were presented as distractors in the Lag-3 position during the RSVP task in the current experiment.

## METHODS

### Participants

Another group of 108 students from the Faculty of Psychology at the University of Barcelona participated in the online version experiment after providing informed consent signed on the same website. We excluded the data from 8 students on the basis of the same criterion outlined in Experiment 1. Consequently, the final sample was composed of 100 participants, who we compensated with course credits for completing the experiment. Fifty-three participants were randomly and automatically assigned to the sweet food condition, whereas the remaining (47) were allocated to the savory food condition upon entering the experiment on the website. The two samples were similar in terms of sex composition, age, and BMI distribution (all *p* values > 0.4; Table S1).

### Stimuli and procedure

All of the stimuli used in the two experimental tasks were identical to those used in Experiment 1. The only difference was that in this experiment, compared with those in Experiment 1, participants were asked to recall the food items immediately after completing the 30-second period provided to memorize them. The RSVP task was administered after memory recall was completed.

## RESULTS

The participants’ performance in Experiment 2 was satisfactorily accurate (95%) when the presence or absence of the target in the sequence without the distractor was identified. Additionally, we observed that task performance was significantly better when the target was absent than when it was presented [target accuracy: *M* = 94.04 ± 5.5; nontarget accuracy: *M* = 96.46 ± 3.20; *F*(1,98) = 14.81; *p* < 0.001; η_p_^2^ = 0.013]. However, we found that neither *food type* [*F*(1,98) = 0.08; *p* = 0.775; η_p_^2^ = 0.008] nor the interaction of *food type* × *target detection* [*F*(1,98) = 2.62; *p* = 0.11; η_p_^2^ = 0.03] reached statistical significance, indicating that the performance of both groups of participants in the RSVP task was similar.

A summary of the results is shown in Figure S1. We began by analyzing the memory performance of the two *food-type* groups. Similar to what we found in Experiment 1, images corresponding to savory foods were easier to memorize than those corresponding to sweet foods [savory foods: *M* = 7.74 ± 0.87; sweet foods: *M* = 6.92 ± 1.12; *t*(98) = 4.10; *p* < 0.001; *d* = 0.81]. The number of distractor images recognized to be seen within the sequences was similar for the two food types [*distractor salience*, savory foods: *M* = 3.66 ± 2.56; sweet foods: *M* = 3.32 ± 2.30; *t*(98) = 0.69; *p* = 0.49; *d* = 0.14], replicating what we observed in Experiment 1.

We subsequently analyzed whether the distractor food images effectively captured the participants’ attention. The results revealed the absence of a significant effect of *Lag* [Lag-3: *M* = 90.59 ± 10.44; Lag-9: *M* = 93.70 ± 5.85; *F*(1,97) = 0.19; *p* = 0.661; η_p_^2^ = 0.002] or *food type* [*F*(1,97) = 0.28; *p* = 0.599; η_p_^2^ = 0.003]. The *Lag* × *food type interaction* was not statistically significant [*F*(1,97) = 1.41; *p* = 0.239; η_p_^2^ = 0.014]. These results indicated that the current experimental setup of food images was ineffective in capturing participants’ attention, as task performance was not affected by food images that appeared in the series in the Lag-3 position, where the EAB is known to occur.

## DISCUSSION

In Experiment 3, we diminished the salience of stimuli in the participants’ mindset by requiring them to complete the assigned task—for which the stimuli were memorized—just before proceeding to the subsequent target detection task. The results indicated that these food stimuli used as distractors were no longer effective in capturing attention after their salience was reduced. Thus, a diminished salience was achieved by rendering them completely irrelevant, as the primary goal task was concluded. The findings from Experiment 3 suggested that an active mindset was necessary for the food images to induce an EAB, as observed in Experiment 1. However, an active mindset with task-relevant stimuli may be necessary but not sufficient for increasing stimuli salience enough to elucidate their attentional capture, as observed in the RSVP in Experiment 1. In other words, even though the stimuli were kept active in Experiment 1 because of the need to complete the incoming memory recall task, they might still need to be directly reexperienced to interfere with the ongoing RSVP task. To explore this possibility, we conducted a new experiment using identical stimuli and procedures as those used in Experiment 1. The only exception in Experiment 4 was that different food items, distinct from those just memorized, were used as distractor images in the RSVP task.

## EXPERIMENT 4

In this experiment, we examined the specificity of stimuli preloaded in participants’ mindsets to induce EAB when those items were still active while completing an incoming task. To further investigate whether the EAB observed toward foods in Experiment 1 was stimuli- specific or could otherwise be generalized to other similar food items, we conducted a new online experiment using the same stimuli and a procedure similar to that in Experiment 1. This time, the participants were instructed to memorize food images from a flyer, but for the subsequent RSVP task, we replaced these food images with those from the other food-type condition. Consequently, with the current experimental setup, we aimed to keep food stimuli active in participants’ mindsets while presenting different images as distractors in the RSVP task. Our goal was to assess whether these novel items were still effective in capturing participants’ attention. We hypothesized that if the EAB is stimulus specific, the new set of food images used as distractors in the RSVP task should not disrupt participants’ performance, as we did in Experiment 1.

## METHODS

### Participants

Another group of 105 students from the Faculty of Psychology at the University of Barcelona participated in the present online experiment. All of the participants provided signed informed consent on the website when they agreed to participate in the experiment. We removed the data from 7 participants on the basis of the exclusion criteria outlined in Experiment 1. The final sample was composed of 98 participants who were randomly and automatically assigned to the savory foods (45) or sweet foods (53) condition at the time of entry into the experiment. The two food conditions were similar in terms of sex, age, and BMI (all *p* values > 0.08) (Table S1). We compensated participants with course credits after the completion of the experiment.

### Stimuli and procedure

We used the same stimuli, tasks, and experimental procedure as in Experiment 1. The only difference was that we orthogonally interchanged the two sets of food types from the memory task to the RSVP task. Thus, participants were still needed, as their main task, to memorize a set of food images presented on a flyer and to recall them at the end of the experiment, exactly as we did in Experiment 1. However, in this case, the food images used as distractors in the RSVP corresponded to different food items than the ones memorized.

## RESULTS

In Experiment 4, mirroring the results observed in all previous experiments, the participants’ overall performance in the RSVP task was highly accurate (95%), with marginally significantly better accuracy in the absence of the target than when it was present [target accuracy: *M* = 94.55 ± 4.85; nontarget accuracy: *M* = 95.76 ± 4.38; *F*(1,96) = 3.73; *p* = 0.056; η_p_^2^ = 0.04]. However, task performance was similar for sweet and savory foods [*food type*: *F*(1,96) = 1.32; *p* = 0.254; η_p_^2^ = 0.01; food *type* × *target detection*: *F*(1,96) < 0.01; *p* = 0.952; η_p_^2^ < 0.001].

The participants’ performance in the memory task indicated that savory food images were again easier to remember than sweet food images were (savory foods: *M* = 6.42 ± 1.78; sweet foods: *M* = 5.70 ± 1.89), although the difference was marginally significant [*t*(96) = 1.95; *p* = 0.054; *d* = 0.39].

We subsequently analyzed whether the food distractors captured the participants’ attention (Figure S1). We did not find an effect of *Lag* [Lag-3: *M* = 93.58 ± 5.96; Lag-9: *M* = 95.31 ± 4.69 *F*(1,95) = 2.35; *p* = 0.129; η_p_^2^ = 0.024] or *food type* [*F*(1,95) = 0.19; *p* = 0.663; η_p_^2^ = 0.002].

Importantly, the interaction term did not reach significance either [*F*(1,95) = 1.18; *p* = 0.281; η_p_^2^ = 0.012]. These results indicate that the enhanced stimulus salience prompted by the current experimental setup was not extensive to other food items, as reflected by the null EAB detected in the current results.

## DISCUSSION

The results obtained in Experiment 4 indicated that preloading food stimuli in memory and keeping them active in participants’ mindsets were not sufficient to increase the salience of other food stimuli to elicit an EAB. This suggests that the EAB observed in Experiment 1 could be defined as stimulus specific. Thus, the momentary capacity acquired by food images to capture attention does not appear to be generalized to other food items, at least under the specific conditions of the RSVP version implemented in the current study.

## ADDITIONAL RESULTS

### BMI distribution across experiments

One-way ANOVA was used to evaluate whether the participants’ BMI was similarly distributed across the experiments. No significant difference emerged among the four experiments [*F*(3,413) = 1.34, *p* = 0.261; see Figure S1].

---

**Figure S1.**
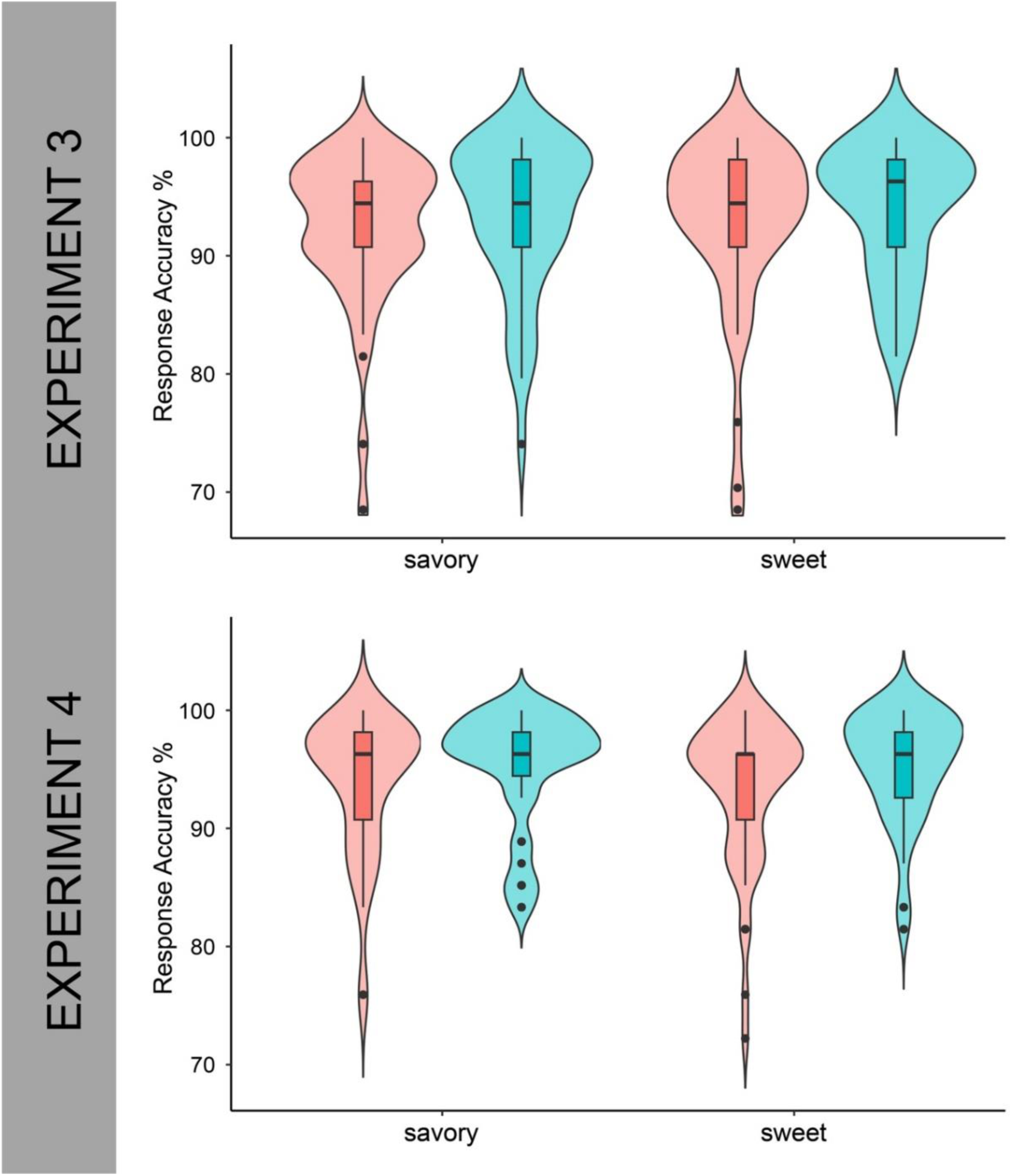
RSVP task performance (Lag 3 vs. Lag 9) in Experiments 3 and 4, separated by food-type condition (savory vs. sweet).

**Figure S2.**
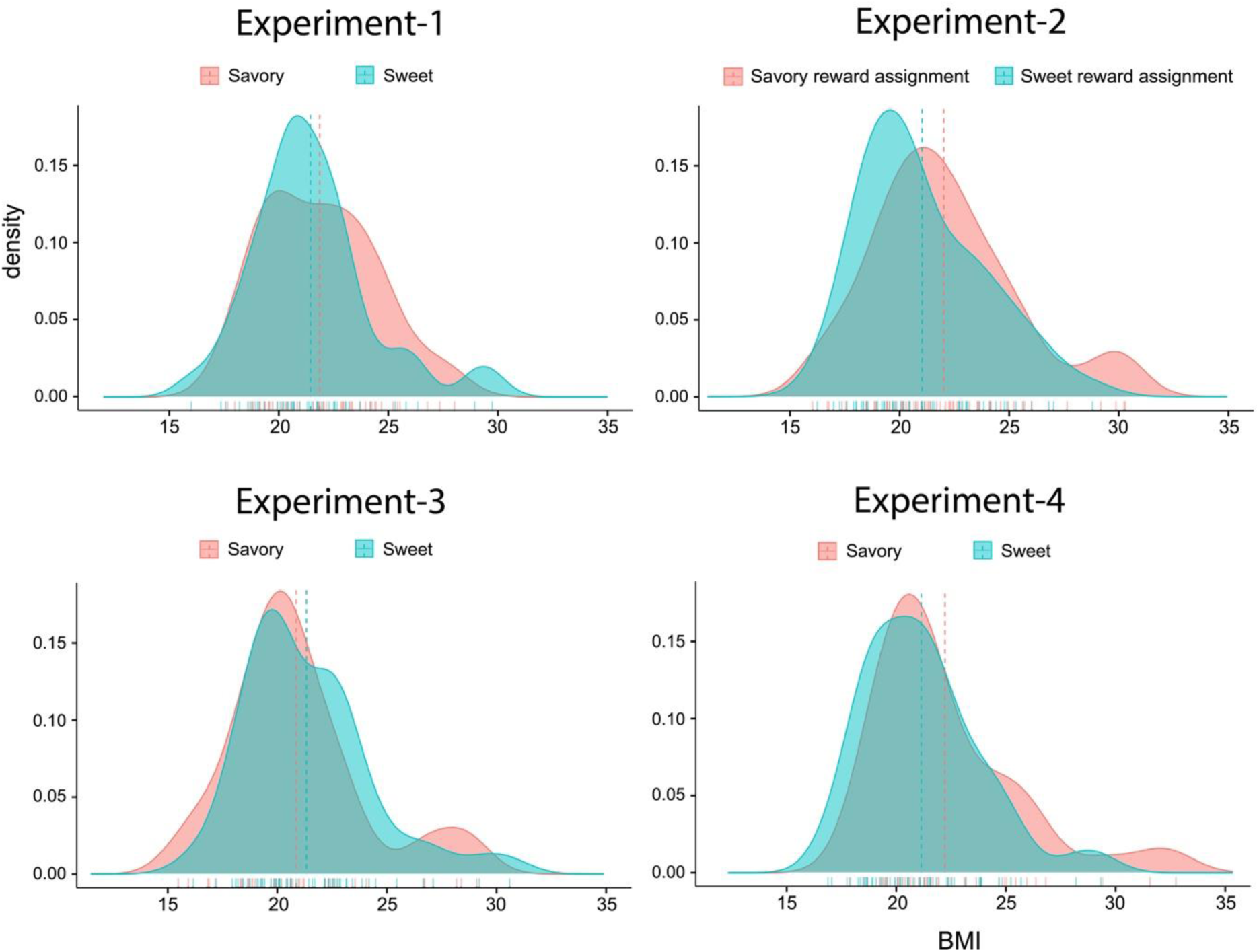
The body mass index (BMI) distribution in all 4 experiments is depicted with density maps. The dashed lines indicate the means in each group (Experiments 1-3: food type: savory vs. sweet; Experiment 4: food-reward assignment: savory-reward assignment vs. sweet-reward assignment).

**Figure S3.**
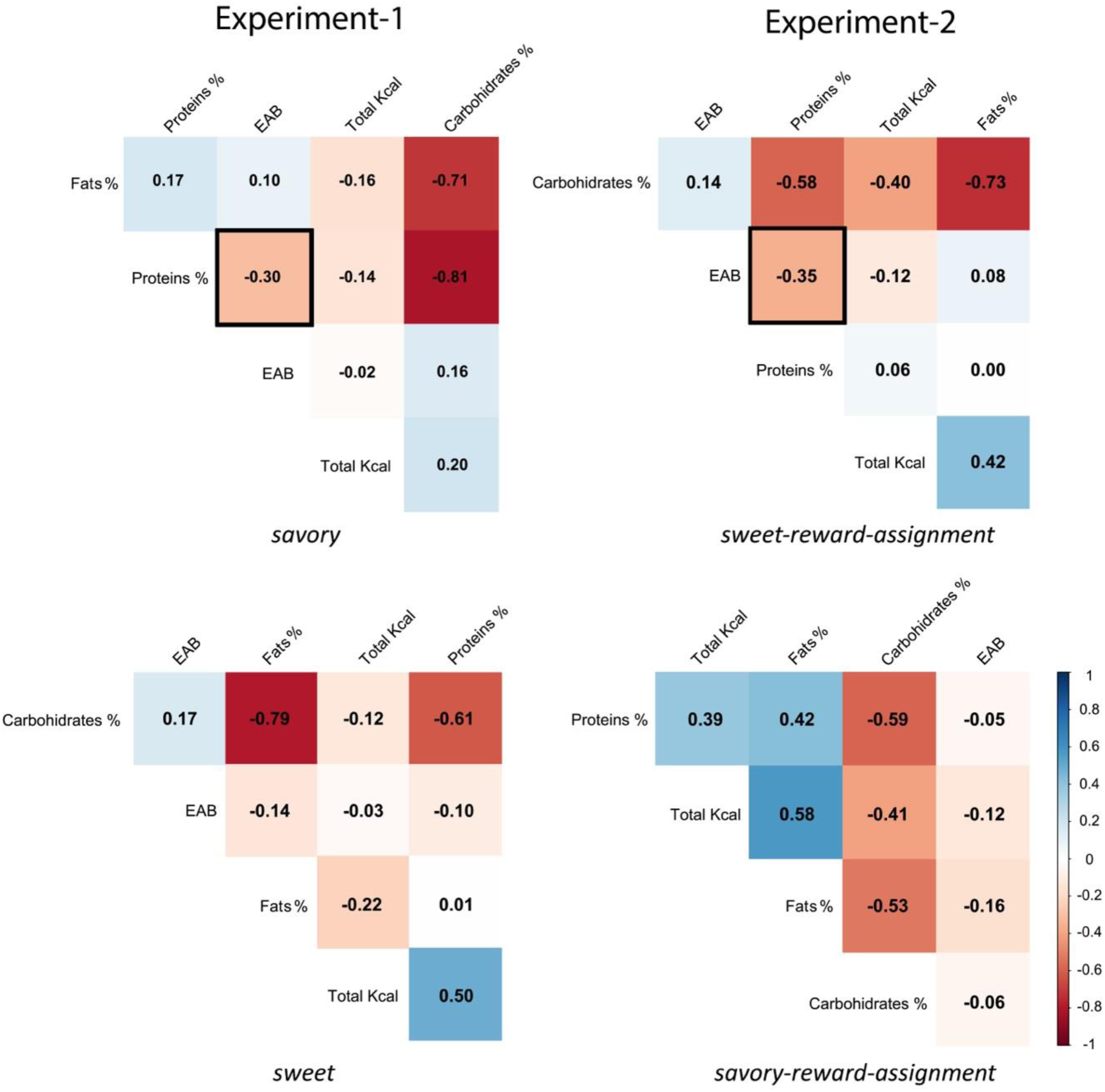
Color heatmaps of Pearson correlations for Experiments 1 and 4. The black frame highlights the sole macronutrient—protein percentage—significantly associated with the EAB effect. The negative values indicate that greater EAB effects occurred with lower protein consumption in participants’ last meal.

**Table S1.**
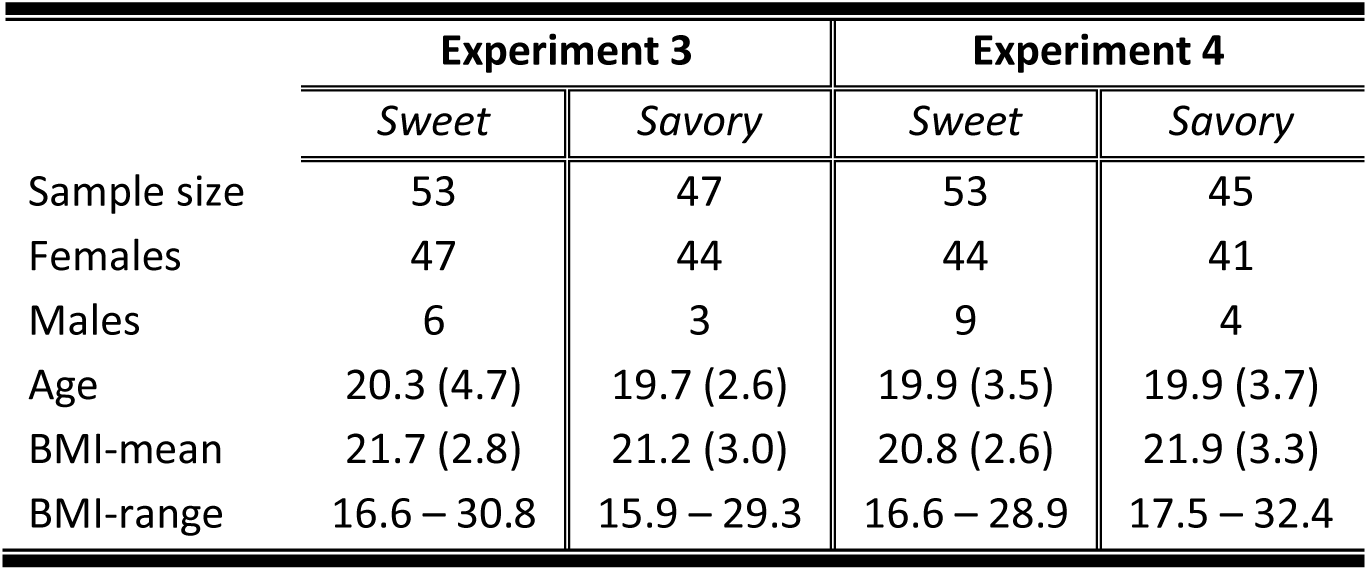
Demographic and anthropometric characteristics of participants in Experiments 3 and 4 by food-type condition (sweet vs. savory). The values are presented as the means with standard deviations in parentheses for age and BMI and as absolute counts for sample size and sex. The BMI range indicates the minimum and maximum body mass index observed within each group.

|  | Experiment 3 |  | Experiment 4 |  |
| --- | --- | --- | --- | --- |
|  | *Sweet* | *Savory* | *Sweet* | *Savory* |
| Sample size | 53 | 47 | 53 | 45 |
| Females | 47 | 44 | 44 | 41 |
| Males | 6 | 3 | 9 | 4 |
| Age | 20.3 (4.7) | 19.7 (2.6) | 19.9 (3.5) | 19.9 (3.7) |
| BMI-mean | 21.7 (2.8) | 21.2 (3.0) | 20.8 (2.6) | 21.9 (3.3) |
| BMI-range | 16.6 – 30.8 | 15.9 – 29.3 | 16.6 – 28.9 | 17.5 – 32.4 |

**Table S2.** Percentage macronutrient composition and total caloric intake for participants in Experiments 1 and 4, shown separately by food-type condition (Experiment 1) and food-reward assignment (Experiment 4). These values were calculated following the methodology detailed in the Methods section, which was validated by a strong interrater correlation (Pearson’s r ≈ 0.8) across all four measures.

| EXPERIMENT-1 | sweet-food | savory-food | sweet vs. savory |  |
| --- | --- | --- | --- | --- |
|  | *Mean (S.D.)* | *Mean (S.D.)* | *t-value (d.f.)* | *p-value* |
| **Total Calories (Kcal)** | 262.16 (172.54) | 303.52 (216.39) | 1.04 (94) | 0.30 |
| **Carbohydrates %** | 64.82 (20.09) | 60.56 (21.86) | 0.99 (94) | 0.32 |
| **Fats %** | 14.95 (13.92) | 16.87 (12.59) | 0.70 (94) | 0.48 |
| **Proteins %** | 20.23 (15.22) | 22.58 (16.98) | 0.71 (94) | 0.47 |
| EXPERIMENT-2 | sweet-reward-assig. | savory-reward-assig. | sweet vs. savory |  |
|  | *Mean (S.D.)* | *Mean (S.D.)* | *t-value (d.f.)* | *p-value* |
| **Total Calories (Kcal)** | 517.95 (257.9) | 514.98 (222.7) | 0.07 (115) | 0.95 |
| **Carbohydrates %** | 45.46 (15.2) | 47.63 (14.6) | -0.79 (115) | 0.43 |
| **Fats %** | 36.45 (12.7) | 34.3 (10.6) | 0.99 (115) | 0.32 |
| **Proteins %** | 18.09 (8.8) | 18.07 (7.5) | 0.01 (115) | 0.99 |

## Footnotes

1 *T* tests comparing participants from the savory and sweet conditions regarding the different macronutrient percentage (*protein*, *carbohydrate*, and *fat*) and total calorie intake in their last meal revealed no differences between groups (all *p values* > .30; see Table S1).

2 We also included in the basic model, one by one, the factors corresponding to the distinctive perceptual properties of foods, and the number of memorized images recalled at the end of the experiment. None of those factors were found to improve the model fit when they were incorporated in the model.

3 Item IDs could not be included as a random effect in the models because they were not recorded in the raw data due to a programming error in the experimental task.

## REFERENCES

Al-Shawaf, L. (2016). The evolutionary psychology of hunger. Appetite, 105, 591–595. 10.1016/j.appet.2016.06.021

Anderson, B. A., Kim, H., Kim, A. J., Liao, M. R., Mrkonja, L., Clement, A., & Grégoire, L. (2021). The past, present, and future of selection history. Neuroscience and Biobehavioral Reviews, 130(April), 326–350. 10.1016/j.neubiorev.2021.09.004

Anderson, B. A., Laurent, P. A., & Yantis, S. (2011). Value-driven attentional capture. Proceedings of the National Academy of Sciences of the United States of America, 108(25), 10367–10371. 10.1073/pnas.1104047108

Arnell, K. M., Killman, K. V., & Fijavz, D. (2007). Blinded by Emotion: Target Misses Follow Attention Capture by Arousing Distractors in RSVP. Emotion, 7(3), 465–477. 10.1037/1528-3542.7.3.465

Arumäe, K., Kreegipuu, K., & Vainik, U. (2019). Assessing the overlap between three measures of food reward. Frontiers in Psychology, 10(APR), 1–11. 10.3389/fpsyg.2019.00883

Ballestero-Arnau, M., Moreno-Sánchez, M., & Cunillera, T. (2021). Food is special by itself: Neither valence, arousal, food appeal, nor caloric content modulate the attentional bias induced by food images. Appetite, 156. 10.1016/j.appet.2020.104984

Bates, D., Mächler, M., Bolker, B., & Walker, S. (2015). Fitting linear mixed-effects models using lme4. Journal of Statistical Software, 67, 1–48. 10.1017/CBO9781107415324.004

Berridge, K. C. (2009a). “Liking” and “wanting” food rewards: Brain substrates and roles in eating disorders. Physiology and Behavior, 97(5), 537–550. 10.1016/j.physbeh.2009.02.044

Berridge, K. C. (2009b). “Liking” and “wanting” food rewards: Brain substrates and roles in eating disorders. Physiology and Behavior, 97(5), 537–550. 10.1016/j.physbeh.2009.02.044

Berthoud, H. R. (2012). The neurobiology of food intake in an obesogenic environment. Proceedings of the Nutrition Society, 71(4), 478–487. 10.1017/S0029665112000602

Berthoud, H. R., Nederkoorn, C., Guerrieri, R., Havermans, R. C., Roefs, A., & Jansen, A. (2007). Interactions between the “cognitive” and “metabolic” brain in the control of food intake. Physiology & Behavior, 91(5), 486–498. 10.1016/j.physbeh.2006.12.016

Birch, L. L. (1999). Development of food preferences. Annual Review of Nutrition, 19(1), 41–62. 10.1146/annurev.nutr.19.1.41

Blechert, J., Lender, A., Polk, S., Busch, N. A., & Ohla, K. (2019). Food-pics_extended-an image database for experimental research on eating and appetite: Additional images, normative ratings and an updated review. Frontiers in Psychology, 10(MAR), 1–9. 10.3389/fpsyg.2019.00307

Cabanac, M. (1971). Physiological Role of Pleasure. Science, 173(4002), 1103–1107. 10.1126/science.173.4002.1103

Cabanac, M., & Fantino, M. (1977). Origin of olfacto-gustatory alliesthesia: Intestinal sensitivity to carbohydrate concentration? Physiology & Behavior, 18(6), 1039–1045. 10.1016/0031-9384(77)90009-9

Campbell, C. P., Raubenheimer, D., Badaloo, A. V., Gluckman, P. D., Martinez, C., Gosby, A., Simpson, S. J., Osmond, C., Boyne, M. S., & Forrester, T. E. (2016). Developmental contributions to macronutrient selection: A randomized controlled trial in adult survivors of malnutrition. *Evolution*, Medicine and Public Health, 2016(1), 158–169. 10.1093/EMPH/EOV030

Castellanos, E. H., Charboneau, E., Dietrich, M. S., Park, S., Bradley, B. P., Mogg, K., & Cowan, R. L. (2009). Obese adults have visual attention bias for food cue images: Evidence for altered reward system function. International Journal of Obesity, 33(9), 1063–1073. 10.1038/ijo.2009.138

Charrondiere, U. R., Chevassus-Agnes, S., Marroni, S., & Burlingame, B. (2004). Impact of different macronutrient definitions and energy conversion factors on energy supply estimations. Journal of Food Composition and Analysis, 17(3–4), 339–360. 10.1016/j.jfca.2004.03.011

Chaumontet, C., Recio, I., Fromentin, G., Benoit, S., Piedcoq, J., Darcel, N., & Tomé, D. (2018). The protein status of rats affects the rewarding value of meals due to their protein content. Journal of Nutrition, 148(6), 989–998. 10.1093/jn/nxy060

Ciesielski, B. G., Armstrong, T., Zald, D. H., & Olatunji, B. O. (2010). Emotion modulation of visual attention: Categorical and temporal characteristics. PLoS ONE, 5(11). 10.1371/journal.pone.0013860

Corbetta, M., & Shulman, G. L. (2002). Control of goal-directed and stimulus-driven attention in the brain. Nature Reviews Neuroscience, 3(3), 201–215. 10.1038/nrn755

Cunningham, C. A., & Egeth, H. E. (2018a). The capture of attention by entirely irrelevant pictures of calorie-dense foods. Psychonomic Bulletin and Review, 25(2), 586–595. 10.3758/s13423-017-1375-8

Cunningham, C. A., & Egeth, H. E. (2018b). The capture of attention by entirely irrelevant pictures of calorie-dense foods. Psychonomic Bulletin and Review, 25(2), 586–595. 10.3758/s13423-017-1375-8

Davidson, G. R., Giesbrecht, T., Thomas, A. M., & Kirkham, T. C. (2018). Pre- and postprandial variation in implicit attention to food images reflects appetite and sensory-specific satiety. Appetite, 125, 24–31. 10.1016/j.appet.2018.01.028

Di Cesare, M., Bentham, J., Stevens, G. A., Zhou, B., Danaei, G., Lu, Y., Bixby, H., Cowan, M. J., Riley, L. M., Hajifathalian, K., Fortunato, L., Taddei, C., Bennett, J. E., Ikeda, N., Khang, Y. H., Kyobutungi, C., Laxmaiah, A., Li, Y., Lin, H. H., … Cisneros, J. Z. (2016). Trends in adult body-mass index in 200 countries from 1975 to 2014: A pooled analysis of 1698 population-based measurement studies with 19.2 million participants. The Lancet, 387(10026), 1377–1396. 10.1016/S0140-6736(16)30054-X

Dobson, K. S., & Dozois, D. J. A. (2004). Attentional biases in eating disorders: A meta-analytic review of Stroop performance. Clinical Psychology Review, 23(8), 1001–1022. 10.1016/j.cpr.2003.09.004

Dux, P. E., & Rentmarois. (2009). The attentional blink: A review of data and theory. *Attention*, Perception, and Psychophysics, 71(8), 1683–1700. 10.3758/APP.71.8.1683

Fiala, N. (2008). Meeting the demand: An estimation of potential future greenhouse gas emissions from meat production. Ecological Economics, 67(3), 412–419. 10.1016/j.ecolecon.2007.12.021

Field, M., Werthmann, J., Franken, I., & Hofmann, W. (2016). The role of attentional bias in obesity and addiction. Health Psychology, 35(8), 767–780. 10.1037/hea0000405

Gosby, A. K., Conigrave, A. D., Lau, N. S., Iglesias, M. A., Hall, R. M., Jebb, S. A., Brand-Miller, J., Caterson, I. D., Raubenheimer, D., & Simpson, S. J. (2011). Testing protein leverage in lean humans: A randomised controlled experimental study. PLoS ONE, 6(10). 10.1371/journal.pone.0025929

Gosby, A. K., Conigrave, A. D., Raubenheimer, D., & Simpson, S. J. (2014). Protein leverage and energy intake. Obesity Reviews, 15(3), 183–191. 10.1111/obr.12131

Greenberg, D., & St. Peter, J. V. (2021). Sugars and sweet taste: Addictive or rewarding? International Journal of Environmental Research and Public Health, 18(18), 1–10. 10.3390/ijerph18189791

Griffioen-Roose, S., Smeets, P. A. M., Van Den Heuvel, E., Boesveldt, S., Finlayson, G., & De Graaf, C. (2014). Human Protein status modulates brain reward responses to food cues 1-3. American Journal of Clinical Nutrition, 100(1), 113–122. 10.3945/ajcn.113.079392

Gutiérrez-Cobo, M. J., Luque, D., Most, S. B., Fernández-Berrocal, P., & Le Pelley, M. E. (2019). Reward and emotion influence attentional bias in rapid serial visual presentation. Quarterly Journal of Experimental Psychology, 72(9), 2155–2167. 10.1177/1747021819840615

Hagan, K. E., Alasmar, A., Exum, A., Chinn, B., & Forbush, K. T. (2020). A systematic review and meta-analysis of attentional bias toward food in individuals with overweight and obesity. Appetite, 151(March), 104710. 10.1016/j.appet.2020.104710

Hall, K. D. (2019). The Potential Role of Protein Leverage in the US Obesity Epidemic. Obesity, 27(8), 1222–1224. 10.1002/oby.22520

Hall, K. D., Ayuketah, A., Brychta, R., Cai, H., Cassimatis, T., Chen, K. Y., Chung, S. T., Costa, E., Courville, A., Darcey, V., Fletcher, L. A., Forde, C. G., Gharib, A. M., Guo, J., Howard, R., Joseph, P. V., McGehee, S., Ouwerkerk, R., Raisinger, K., … Zhou, M. (2019). Ultra-Processed Diets Cause Excess Calorie Intake and Weight Gain: An Inpatient Randomized Controlled Trial of Ad Libitum Food Intake. Cell Metabolism, 30(1), 67–77.e3. 10.1016/j.cmet.2019.05.008

Hardman, C. A., Jones, A., Burton, S., Duckworth, J. J., McGale, L. S., Mead, B. R., Roberts, C. A., Field, M., & Werthmann, J. (2021). Food-related attentional bias and its associations with appetitive motivation and body weight: A systematic review and meta-analysis. Appetite, 157(October 2020), 104986. 10.1016/j.appet.2020.104986

Hendrikse, J. J., Cachia, R. L., Kothe, E. J., Mcphie, S., Skouteris, H., & Hayden, M. J. (2015). Attentional biases for food cues in overweight and individuals with obesity: A systematic review of the literature. Obesity Reviews, 16(5), 424–432. 10.1111/obr.12265

Higgs, S., Robinson, E., & Lee, M. (2012). Learning and Memory Processes and Their Role in Eating: Implications for Limiting Food Intake in Overeaters. Current Obesity Reports, 1(2), 91–98. 10.1007/s13679-012-0008-9

Higgs, S., Rutters, F., Thomas, J. M., Naish, K., & Humphreys, G. W. (2012). Top down modulation of attention to food cues via working memory. Appetite, 59(1), 71–75. 10.1016/j.appet.2012.03.018

Hill, J. O., & Peters, J. C. (1998). Environmental Contributions to the Obesity Epidemic. Science, 280(June), 1371–1374.

Holkner, A., & Pyglet-developers. (2021). Pyglet: A cross-platform windowing and multimedia library for Python.

Jensen, K., Mayntz, D., Toft, S., Clissold, F. J., Hunt, J., Raubenheimer, D., & Simpson, S. J. (2012). Optimal foraging for specific nutrients in predatory beetles. Proceedings of the Royal Society B: Biological Sciences, 279(1736), 2212–2218. 10.1098/rspb.2011.2410

Jiang, T., Soussignan, R., Rigaud, D., Martin, S., Royet, J. P., Brondel, L., & Schaal, B. (2008). Alliesthesia to food cues: Heterogeneity across stimuli and sensory modalities. Physiology and Behavior, 95(3), 464–470. 10.1016/j.physbeh.2008.07.014

Johnson, A. W. (2013). Eating beyond metabolic need: How environmental cues influence feeding behavior. Trends in Neurosciences, 36(2), 101–109. 10.1016/j.tins.2013.01.002

Kaisari, P., Kumar, S., Hattersley, J., Dourish, C. T., Rotshtein, P., & Higgs, S. (2019). Top-down guidance of attention to food cues is enhanced in individuals with overweight/obesity and predicts change in weight at one-year follow up. International Journal of Obesity, 43(9), 1849– 1858. 10.1038/s41366-018-0246-3

Kellett, G. L., Brot-Laroche, E., Mace, O. J., & Leturque, A. (2008). Sugar absorption in the intestine: The role of GLUT2. Annual Review of Nutrition, 28, 35–54. 10.1146/annurev.nutr.28.061807.155518

Kennedy, B. L., Camara, A. M., & Tran, D. M. D. (2024). You eye what you eat: BMI, consumption patterns, and dieting status predict temporal attentional bias to food-associated images. Appetite, 192(June 2023), 107095. 10.1016/j.appet.2023.107095

Kennedy, B. L., & Most, S. B. (2015). Affective stimuli capture attention regardless of categorical distinctiveness: An emotion-induced blindness study. Visual Cognition, 23(1–2), 105–117. 10.1080/13506285.2015.1024300

Kirsten, H., Seib-Pfeifer, L. E., Koppehele-Gossel, J., & Gibbons, H. (2019). Food has the right of way: Evidence for prioritised processing of visual food stimuli irrespective of eating style. Appetite, 142(June), 104372. 10.1016/j.appet.2019.104372

Kohanmoo, A., Faghih, S., & Akhlaghi, M. (2020). Effect of short- and long-term protein consumption on appetite and appetite-regulating gastrointestinal hormones, a systematic review and meta-analysis of randomized controlled trials. Physiology and Behavior, 226(August), 113123. 10.1016/j.physbeh.2020.113123

Lang, P., Bradley, M., & Cuthbert, B. (1997). Motivated attention: affect, activation, and action. In R. F. Simons, P. J. Lang, & M. Balaban (Eds.), Attention and orienting: sensory and motivational processes (pp. 97–136). Lawrence Erlbaum Associates, Inc.

Leidy, H. J., Lepping, R. J., Savage, C. R., & Harris, C. T. (2011). Neural responses to visual food stimuli after a normal vs. higher protein breakfast in breakfast-skipping teens: A pilot fmri study. Obesity, 19(10), 2019–2025. 10.1038/oby.2011.108

Le Pelley, M. E., Mitchell, C. J., Beesley, T., George, D. N., & Wills, A. J. (2016). Attention and associative learning in humans: An integrative review. Psychological Bulletin, 142(10), 1111– 1140. 10.1037/bul0000064

Lieberman, L. S. (2006). Evolutionary and anthropological perspectives on optimal foraging in obesogenic environments. Appetite, 47(1), 3–9. 10.1016/j.appet.2006.02.011

Liu, S., Globa, A. K., Mills, F., Naef, L., Qiao, M., Bamji, S. X., & Borgland, S. L. (2016). Consumption of palatable food primes food approach behavior by rapidly increasing synaptic density in the VTA. Proceedings of the National Academy of Sciences, 113(9), 2520–2525. 10.1073/pnas.1515724113

Liu, Y., Roefs, A., Werthmann, J., & Nederkoorn, C. (2019). Dynamics of attentional bias for food in adults, children, and restrained eaters. Appetite, 135(201706990024), 86–92. 10.1016/j.appet.2019.01.004

Martinez-Cordero, C., Kuzawa, C. W., Sloboda, D. M., Stewart, J., Simpson, S. J., & Raubenheimer, D. (2012). Testing the Protein Leverage Hypothesis in a free-living human population. Appetite, 59(2), 312–315. 10.1016/j.appet.2012.05.013

Martínez Steele, E., Raubenheimer, D., Simpson, S. J., Baraldi, L. G., & Monteiro, C. A. (2018). Ultra-processed foods, protein leverage and energy intake in the USA. Public Health Nutrition, 21(1), 114–124. 10.1017/S1368980017001574

Marty, L., Nicklaus, S., Miguet, M., Chambaron, S., & Monnery-Patris, S. (2018). When do healthiness and liking drive children’s food choices? The influence of social context and weight status. Appetite, 125(June), 466–473. 10.1016/j.appet.2018.03.003

McHugo, M., Olatunji, B. O., & Zald, D. H. (2013). The emotional attentional blink: What we know so far. Running title: The emotional attentional blink. Frontiers in Human Neuroscience, 7(APR 2013), 1–9. 10.3389/fnhum.2013.00151

Most, S. B., Chun, M. M., Widders, D. M., Haven, N., & Zald, D. H. (2005). Attentional rubbernecking: Cognitive control and personality in emotion-induced blindness. Psychonomic Bulletin & Review, 12(4), 654–661.

Murphy, M., Peters, K. Z., Denton, B. S., Lee, K. A., Chadchankar, H., & McCutcheon, J. E. (2018). Restriction of dietary protein leads to conditioned protein preference and elevated palatability of protein-containing food in rats. Physiology and Behavior, 184(December 2017), 235–241. 10.1016/j.physbeh.2017.12.011

Neal, C., Pepper, G. V., Allen, C., & Nettle, D. (2023). No effect of hunger on attentional capture by food cues: Two replication studies. Appetite, 191(May), 107065. 10.1016/j.appet.2023.107065

Neimeijer, R. A. M., de Jong, P. J., & Roefs, A. (2013a). Temporal attention for visual food stimuli in restrained eaters. Appetite, 64, 5–11. 10.1016/j.appet.2012.12.013

Neimeijer, R. A. M., de Jong, P. J., & Roefs, A. (2013b). Temporal attention for visual food stimuli in restrained eaters. Appetite, 64, 5–11. 10.1016/j.appet.2012.12.013

Nijs, I. M. T., Muris, P., Euser, A. S., & Franken, I. H. A. (2010). Differences in attention to food and food intake between overweight/obese and normal-weight females under conditions of hunger and satiety. Appetite, 54(2), 243–254. 10.1016/j.appet.2009.11.004

Parker, R. W. R., Blanchard, J. L., Gardner, C., Green, B. S., Hartmann, K., Tyedmers, P. H., & Watson, R. A. (2018). Fuel use and greenhouse gas emissions of world fisheries. Nature Climate Change, 8(4), 333–337. 10.1038/s41558-018-0117-x

Pearson, D., Watson, P., Albertella, L., & Le Pelley, M. E. (2022). Attentional economics links value-modulated attentional capture and decision-making. Nature Reviews Psychology, 1(6), 320–333. 10.1038/s44159-022-00053-z

Piech, R. M., Pastorino, M. T., & Zald, D. H. (2010). All I saw was the cake. Hunger effects on attentional capture by visual food cues. Appetite, 54(3), 579–582. 10.1016/j.appet.2009.11.003

Plailly, J., Luangraj, N., Nicklaus, S., Issanchou, S., Royet, J. P., & Sulmont-Rossé, C. (2011). Alliesthesia is greater for odors of fatty foods than of non-fat foods. Appetite, 57(3), 615–622. 10.1016/j.appet.2011.07.006

Rangel, A. (2013). Regulation of dietary choice by the decision-making circuitry. Nature Neuroscience, 16(12), 1717–1724. 10.1038/nn.3561

Raubenheimer, D., & Simpson, S. J. (2019). Protein Leverage: Theoretical Foundations and Ten Points of Clarification. Obesity, 27(8), 1225–1238. 10.1002/oby.22531

Raymond, J. E., Shapiro, K. L., & Arnell, K. M. (1992). Temporary Suppression of Visual Processing in an RSVP Task: An Attentional Blink? Journal of Experimental Psychology: Human Perception and Performance, 18(3), 849–860. 10.1037/0096-1523.18.3.849

Richards, M. P., & Schmitz, R. W. (2008). Isotope evidence for the diet of the Neanderthal type specimen. Antiquity, 82(317), 553–559. 10.1017/S0003598X00097210

Richards, M. P., & Trinkaus, E. (2009). Isotopic evidence for the diets of European Neanderthals and early modern humans. Proceedings of the National Academy of Sciences of the United States of America, 106(38), 16034–16039. 10.1073/pnas.0903821106

Roden, M., & Bernroider, E. (2003). Hepatic glucose metabolism in humans - Its role in health and disease. Best Practice and Research: Clinical Endocrinology and Metabolism, 17(3), 365–383. 10.1016/S1521-690X(03)00031-9

Rutters, F., Kumar, S., Higgs, S., & Humphreys, G. W. (2015). Electrophysiological evidence for enhanced representation of food stimuli in working memory. Experimental Brain Research, 233(2), 519–528. 10.1007/s00221-014-4132-5

Santacroce, L. A., Swami, A. L., & Tamber-Rosenau, B. J. (2023). More than a feeling: The emotional attentional blink relies on non-emotional “pop out,” but is weak compared to the attentional blink. *Attention*, Perception, and Psychophysics, 85(4), 1034–1053. 10.3758/s13414-023-02677-6

Schwabe, L., Merz, C. J., Walter, B., Vaitl, D., Wolf, O. T., & Stark, R. (2011). Emotional modulation of the attentional blink: The neural structures involved in capturing and holding attention. Neuropsychologia, 49(3), 416–425. 10.1016/j.neuropsychologia.2010.12.037

Seeley, R. J., & Berthoud, H. R. (2000). Neural and metabolic control of macronutrient selection: consensus and controversy. Berthoud HR, Seeley RJ, Eitors. Neural and Metabolic Control of Macronutrient Intake. CRC Press: Boca Raton, 489–496.

Seibt, B., Häfner, M., & Deutsch, R. (2007). Prepared to eat: How immediate affective and motivational responses to food cues are influenced by food deprivation. European Journal of Social Psychology, 37(2), 359–379. 10.1002/ejsp.365

Shapiro, K. L., Arnell, K. M., & Raymond, J. E. (1997). The attentional blink. Trends in Cognitive Sciences, 1(8), 291–296. 10.1016/S1364-6613(97)01094-2

Simpson, S. J., Batley, R., & Raubenheimer, D. (2003). Geometric analysis of macronutrient intake in humans: The power of protein? Appetite, 41(2), 123–140. 10.1016/S0195-6663(03)00049-7

Simpson, S. J., & Raubenheimer, D. (2005). Obesity: The protein leverage hypothesis. Obesity Reviews, 6(2), 133–142. 10.1111/j.1467-789X.2005.00178.x

Simpson, S. J., & Raubenheimer, D. (2014a). Processed foods that dilute protein content subvert our appetite control. Nature, 191, 2014.

Simpson, S. J., & Raubenheimer, D. (2014b). Processed foods that dilute protein content subvert our appetite control. Nature, 191, 2014.

Simpson, S. J., Sibly, R. M., Lee, K. P., Behmer, S. T., & Raubenheimer, D. (2004). Optimal foraging when regulating intake of multiple nutrients. Animal Behaviour, 68(6), 1299–1311. 10.1016/j.anbehav.2004.03.003

Southgate, D. A. T., & Durnin, J. V. G. A. (1970). Calorie conversion factors. An experimental reassessment of the factors used in the calculation of the energy value of human diets. British Journal of Nutrition, 24(2), 517–535. 10.1079/bjn19700050

Stoet, G. (2017). PsyToolkit: A Novel Web-Based Method for Running Online Questionnaires and Reaction-Time Experiments. Teaching of Psychology, 44(1), 24–31. 10.1177/0098628316677643

Swinburn, B. A., Sacks, G., Hall, K. D., McPherson, K., Finegood, D. T., Moodie, M. L., & Gortmaker, S. L. (2011). The global obesity pandemic: Shaped by global drivers and local environments. The Lancet, 378(9793), 804–814. 10.1016/S0140-6736(11)60813-1

Theeuwes, J. (2019). Goal-driven, stimulus-driven, and history-driven selection. Current Opinion in Psychology, 29, 97–101. 10.1016/j.copsyc.2018.12.024

van Rossum, C. T. M., Fransen, H. P., Verkaik-Kloosterman, J., Buurma-Rethans, E. J. M., & Ocké, M. C. (2011). Dutch National Food Consumption Survey 2007-2010: Diet of children and adults aged 7 to 69 years. Rijksinstituut voor Volksgezondheid en Milieu RIVM.

Werthmann, J., Jansen, A., & Roefs, A. (2016). Make up your mind about food: A healthy mindset attenuates attention for high-calorie food in restrained eaters. Appetite, 105, 53–59. 10.1016/j.appet.2016.05.005

WHO. (2016). Obesity and overweight: Fact sheet 311. WHO Media Centre. Retrieved from http://www.who.int/mediacentre/factsheets/fs311/en/

Wu, G. (2009). Amino acids: Metabolism, functions, and nutrition. Amino Acids, 37(1), 1–17. 10.1007/s00726-009-0269-0

Yokum, S., Ng, J., & Stice, E. (2011). Attentional bias to food images associated with elevated weight and future weight gain: An fMRI study. Obesity, 19(9), 1775–1783. 10.1038/oby.2011.168

Zheng, H., & Berthoud, H. R. (2008). Neural systems controlling the drive to eat: Mind versus metabolism. Physiology, 23(2), 75–83. 10.1152/physiol.00047.2007

